# HIV proviral burden, genetic diversity, and dynamics in viremic controllers who subsequently initiated suppressive antiretroviral therapy

**DOI:** 10.1101/2021.03.29.437632

**Authors:** F. Harrison Omondi, Hanwei Sudderuddin, Aniqa Shahid, Natalie N. Kinloch, Bradley R. Jones, Rachel L. Miller, Olivia Tsai, Daniel MacMillan, Alicja Trocha, Mark A. Brockman, Chanson J. Brumme, Jeffrey B. Joy, Richard Liang, Bruce D. Walker, Zabrina L. Brumme

## Abstract

Curing HIV will require eliminating the reservoir of integrated, replication-competent proviruses that persist despite antiretroviral therapy (ART). Understanding the burden, genetic diversity and longevity of persisting proviruses in diverse individuals with HIV is critical to this goal, but these characteristics remain understudied in some groups. Among them are viremic controllers, individuals who naturally suppress HIV to low levels but for whom therapy is nevertheless recommended. We reconstructed within-host HIV evolutionary histories from longitudinal single-genome amplified viral sequences in four viremic controllers who eventually initiated ART, and used this information to characterize the age and diversity of proviruses persisting on therapy. We further leveraged these within-host proviral age distributions to estimate rates of proviral turnover prior to ART. This is an important yet understudied metric, since pre-ART proviral turnover dictates reservoir composition at ART initiation (and thereafter), which is when curative interventions, once developed, would be administered. Despite natural viremic control, all participants displayed significant within-host HIV evolution pre-therapy, where overall on-ART proviral burden and diversity broadly reflected the extent of viral replication and diversity pre-ART. Consistent with recent studies of non-controllers, the proviral pools of two participants were skewed towards sequences that integrated near ART initiation, suggesting dynamic proviral turnover during untreated infection. In contrast, proviruses recovered from the two other participants dated to time-points that were more evenly spread throughout infection, suggesting slow or negligible proviral decay following deposition. HIV cure strategies will need to overcome within-host proviral diversity, even in individuals who naturally controlled HIV replication before therapy.

**Importance:** HIV therapy is life-long because integrated, replication-competent viral copies persist within long-lived cells. To cure HIV, we need to understand when these viral reservoirs form, how large and genetically diverse they are, and how long they endure. Elite controllers, individuals who naturally suppress HIV to undetectable levels, are being intensely studied as models of HIV remission, but viremic controllers, individuals who naturally suppress HIV to low levels, remain understudied even though they too may hold valuable insights. We combined phylogenetics and mathematical modeling to reconstruct proviral seeding and decay from infection to therapy-mediated suppression in four viremic controllers. We recovered diverse proviruses persisting during therapy that broadly reflected HIV’s within-host evolutionary history, where the estimated half-lives of the persistent proviral pool during untreated infection ranged from <1 year to negligible. Cure strategies will need to contend with proviral diversity and between-host heterogeneity, even in individuals who naturally control HIV.

## Background

Like all retroviruses, HIV integrates its genome into that of its host cell. Most infected cells die − or are eliminated by the immune system − within a day or two of infection, but a small number persist long-term, even during suppressive antiretroviral therapy (ART) (1–3). While most of these long-lived cells harbor HIV proviruses with large deletions or other genetic defects (4–6), a minority harbor replication-competent proviruses that could reactivate at any time. It is for this reason that ART needs to be maintained for life. Achieving ART-free HIV remission (“functional cure”) or a complete cure (eradication) (7) will therefore require permanently inactivating (8, 9) or eliminating (10, 11) these HIV reservoirs, respectively. Characterizing the burden, genetic diversity and longevity of the proviruses that persist within-host can advance us towards these goals.

HIV elite controllers, rare individuals who spontaneously control viremia to undetectable levels without ART, are being intensely studied as natural models of HIV remission (12). They may also represent a group in which a complete cure might be more easily achieved (13, 14), in part because their HIV reservoirs are smaller and less genetically diverse (8, 12, 15). Proviral burden and diversity however are less well characterized in viremic controllers, individuals who naturally suppress HIV to low levels (normally defined as <2000 HIV RNA copies/ml in plasma (16, 17)), but who are nevertheless recommended to initiate ART under current guidelines (18). In particular, this group could yield insights into persisting proviral composition in the context of natural yet incomplete HIV control.

More broadly, our understanding of within-host proviral *dynamics* (*i.e.* the rates of proviral deposition and subsequent decay) during untreated infection remains incomplete for HIV controllers in general. This is in part because longitudinal studies dating back to infection are rare in this group, and because it is challenging to isolate HIV from samples with low or undetectable viral loads. Understanding these dynamics is nevertheless important because proviral longevity during untreated infection dictates reservoir composition at ART initiation (and thereafter): if the persisting proviral pool turned over slowly pre-ART, then HIV sequences seeded into it during early infection would have a high likelihood of persisting for long periods, but if the pool turned over rapidly, its composition would shift towards recently-circulating HIV sequences. As cure or remission interventions would only be administered during suppressive ART, it is critical to understand the dynamics that shape the proviral pool up to this point.

Our understanding of within-host proviral dynamics has recently been enriched by the development of phylogenetic approaches to analyze persisting proviral diversity in context of HIV’s within-host evolution prior to ART (19–23). A recent study that recovered replication-competent HIV sequences during therapy revealed that the majority represented viral variants that circulated close to therapy initiation, though more ancestral sequences, some dating back to transmission, were also recovered (21). Studies employing similar approaches to “date” the overall pool of proviruses persisting on ART, which comprise both genetically intact and defective sequences, have confirmed that these often span the individual’s whole pre-ART history, where individuals differ in the extent to which their proviral pool is skewed towards sequences that integrated around ART initiation (19, 20, 22, 23). Because viremic controllers naturally limit viral replication to low levels, one could hypothesize that their persisting proviral pools may be smaller and less diverse than those of non-controllers, which could potentially make them more responsive to remission or cure interventions, but on-ART proviral diversity has not been investigated in this group in context of HIV’s full within-host evolutionary history.

The age distributions of proviruses sampled shortly after ART can also be used to calculate within-host rates of proviral turnover during untreated infection (19, 23): this is because, at the time of sampling, the majority of proviral turnover would have already occurred prior to therapy. Recent data from non-controllers suggests that proviral half-lives during untreated infection average less than a year (19, 24), though another study estimated an average of 25 months (23). Regardless, these estimates are far shorter than the published rates of proviral decay on ART, which are estimated as ∼4 years for the replication competent reservoir (25) and >10 years for the proviral pool at large (26). This has led to the hypothesis that ART dramatically slows the rate of proviral turnover *in vivo*, thereby stabilizing the proviral pool for long-term persistence (21, 27, 28). Pre-ART proviral dynamics however have not been investigated in HIV controllers. To address these knowledge gaps, we combined single-genome sequencing, proviral quantification, phylogenetic analysis, and mathematical modeling to characterize proviral burden, diversity and dynamics in four viremic controllers from HIV infection to therapy-mediated suppression.

## Results

### Participant characteristics and sampling

We studied four viremic controllers. Three broadly maintained plasma viral loads (pVL) <2000 copies HIV RNA/ml during untreated infection (participants 1, 2, 3), while one maintained pVL <2000 copies HIV RNA/ml for 3 years before losing control (participant 4). Participants initiated ART a mean of 6.1 (range 3.8-8.4) years following their estimated date of infection (EDI). An average of 13 (range 10-17) pre-ART plasma samples per participant, that spanned a mean 4.5 (range 2.5-6.7) year period prior to therapy, were available for HIV RNA sequencing, where the first was sampled a mean of 20.5 (range 15-26) months following the EDI (Figure 1; Table 1). In addition, PBMC were available at two timepoints on suppressive ART (10 million cells per timepoint). Due to limited cell numbers, the first PBMC sample, taken an average of 18.7 (range 9-38) months after ART, was used to quantify total and genomically intact proviral burden in CD4+ T cells (29). The second, taken an average of 28.8 (range 16-54) months after ART, was used for proviral sequencing and integration date inference (20).

**Fig. 1:**
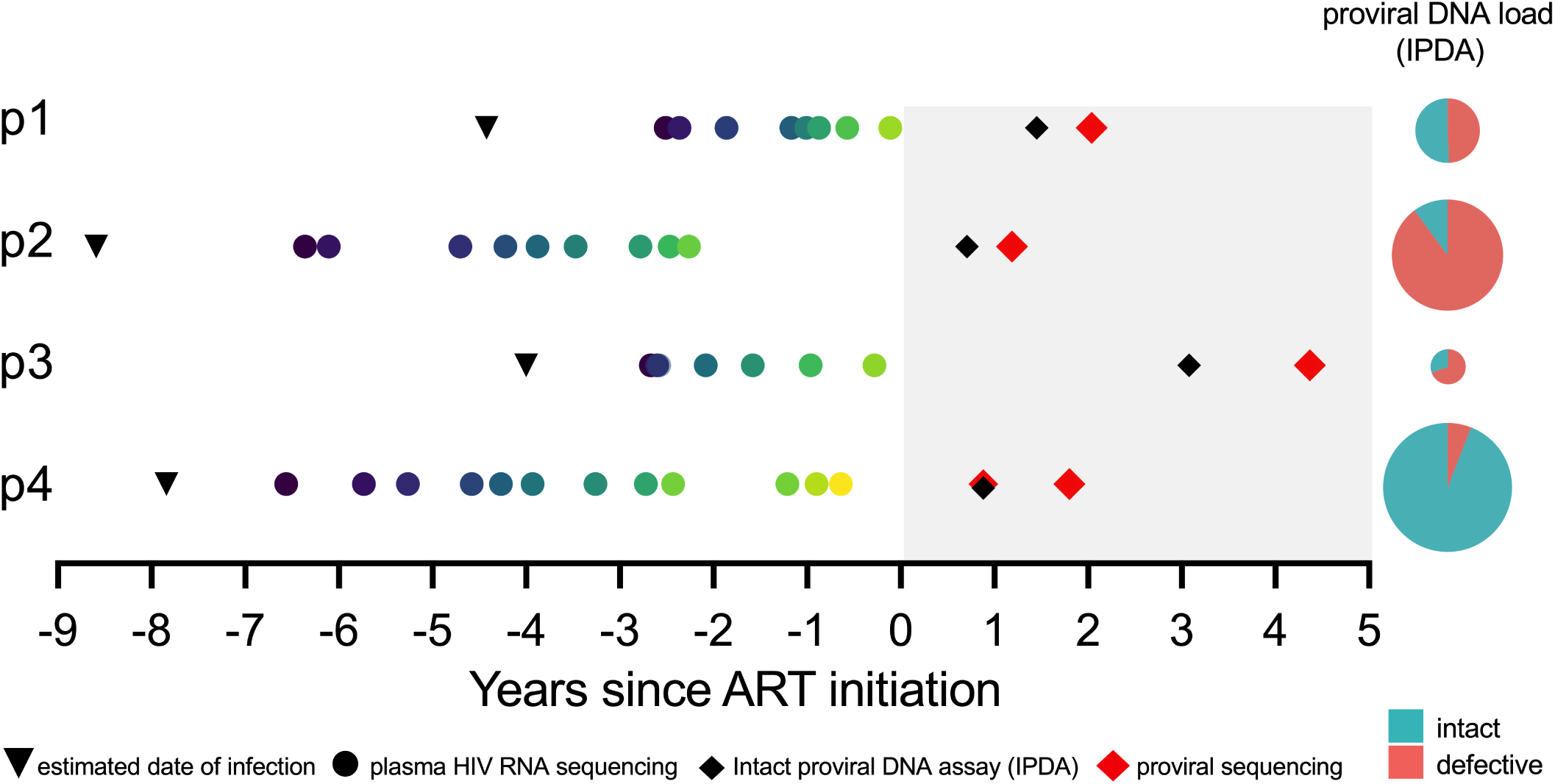
Participant sampling timeline and reservoir quantification. Timeline is depicted as years since ART initiation. Shading represents ART. Inverted black triangles denote the clinically-estimated date of infection. Coloured circles denote pre-ART plasma samples from which HIV RNA sequences were isolated. Red diamonds denote cell samples on ART from which proviral sequences were isolated. Black diamonds denote proviral quantification dates using the Intact Proviral DNA Assay (IPDA). IPDA results are shown as pie charts, where the pie size denotes the total proviral burden and the coloured slices denote intact and defective HIV genome proportions (actual values shown in Table 1).

**Table 1:** Participant clinical and HIV sequence sampling details

| Participant | Clinically estimated infection date | Diagnosis date | First plasma viral load<sup>a</sup> | N Plasma timepoints with HIV sequence data<sup>b</sup> | N Plasma HIV sequences (distinct: N; %)<sup>c</sup> | ART initiation date | Total proviral burden on ART (% genomically intact)<sup>d</sup> | Proviral sequencing date(s) | N Proviral sequences (distinct: N, %)<sup>c</sup> |
| --- | --- | --- | --- | --- | --- | --- | --- | --- | --- |
| 1 | Apr 2006 | Feb 2007 | 1615 | 8 | 104 (52; 50%) | 2010-08-01 | 215 (51) | 2012-09-13 | 48 (40, 83%) |
| 2 | Dec 2005 | Jan 2006 | 977 | 9 | 67 (60; 90%) | 2014-05-01 | 580 (10) | 2015-09-08 | 66 (36, 55%) |
| 3 | Mar 2008 | Jun 2008 | 97 | 7 | 130 (71; 55%) | 2012-01-01 | 44 (31) | 2016-07-14 | 24 (12, 50%) |
| 4 | Mar 2005 | Nov 2005 | 383 | 12 | 245 (173; 71%) | 2013-02-01 | 778 (94) | 2013-11-18;<br>2014-10-20 | 1 full genome<br>128 (119, 92%) |

<sup>a</sup> First detectable sampled plasma viral load, expressed as HIV RNA copies/ml plasma
<sup>b</sup> Total number of plasma timepoints from which HIV RNA *nef* sequences were successfully amplified.
<sup>c</sup> "Distinct" refers to sequences that were observed only once in that compartment. Reported Ns exclude defective, hypermutated and putative within-host recombinant HIV sequences.
<sup>d</sup> as measured by the Intact Proviral DNA Assay and expressed as HIV copies/10<sup>6</sup> CD4+ T cells.

### Quantifying total and genetically intact proviral burden

We quantified total and genomically intact proviral burden in CD4+ T cells using the Intact Proviral DNA Assay (IPDA). This is a duplexed droplet digital PCR (ddPCR) assay that targets two HIV regions, the Packaging Signal (Ψ) near the 5’ end of the viral genome and the Rev Responsive Element (RRE) within Envelope (*env*) that, together, have high predictive power to distinguish genomically intact proviruses from those with large deletions and/or extensive hypermutation, which typically dominate *in vivo* (29). Proviruses yielding double (Ψ and *env*) positive signals are inferred to be genetically intact while those yielding only single positive signal are classified as defective.

We observed marked inter-individual differences in total proviral burden. Participant 3 harbored the fewest proviruses (44 HIV DNA copies/10^6^ CD4+ T cells) while participant 4, the individual who eventually lost control, harbored the most (778 HIV DNA copies/10^6^ CD4+ T cells) (Figure 1; Table 1). The percent of genomically intact proviruses also differed markedly, with participant 2 harboring the lowest (10% intact proviruses) and participant 4 the highest (94% intact). The latter is remarkable though not without precedent (30). Participant 4’s large overall and intact proviral burden allowed us to isolate one near-full-length intact provirus from this timepoint (Figure 1) for integration date inference.

### Single-genome plasma HIV RNA and proviral sequencing

We characterized HIV RNA *nef* sequences from longitudinal pre-ART plasma, and proviral sequences from the second on-ART PBMC sample, by single-genome amplification. HIV *nef* was sequenced because of its richness in phylogenetic signal despite its relatively short length (a shorter amplicon was critical as viral loads were low for most samples), because *nef* is representative of within-host HIV diversity and evolution relative to the rest of the viral genome (20), and because it is the gene most likely to be intact in proviruses persisting on ART (29, 31). Despite low viral loads we isolated 546 HIV RNA *nef* sequences from pre-ART plasma (range 67-245/participant), where an average of 66.2% of these (range 50.0-89.6%) were distinct within each participant (Table 1). We further isolated 267 intact proviral *nef* sequences during suppressive ART (range 24-129/participant), where an average of 69.8% (range 50.0-91.5%) were distinct. All participants had HIV subtype B and their sequences were monophyletic with no evidence of co-infection or superinfection (Figure 2). There were multiple instances where we recovered a proviral sequence on ART that was identical to a pre-ART plasma sequence, consistent with the long-term persistence of HIV sequences *in vivo*.

**Fig. 2:**
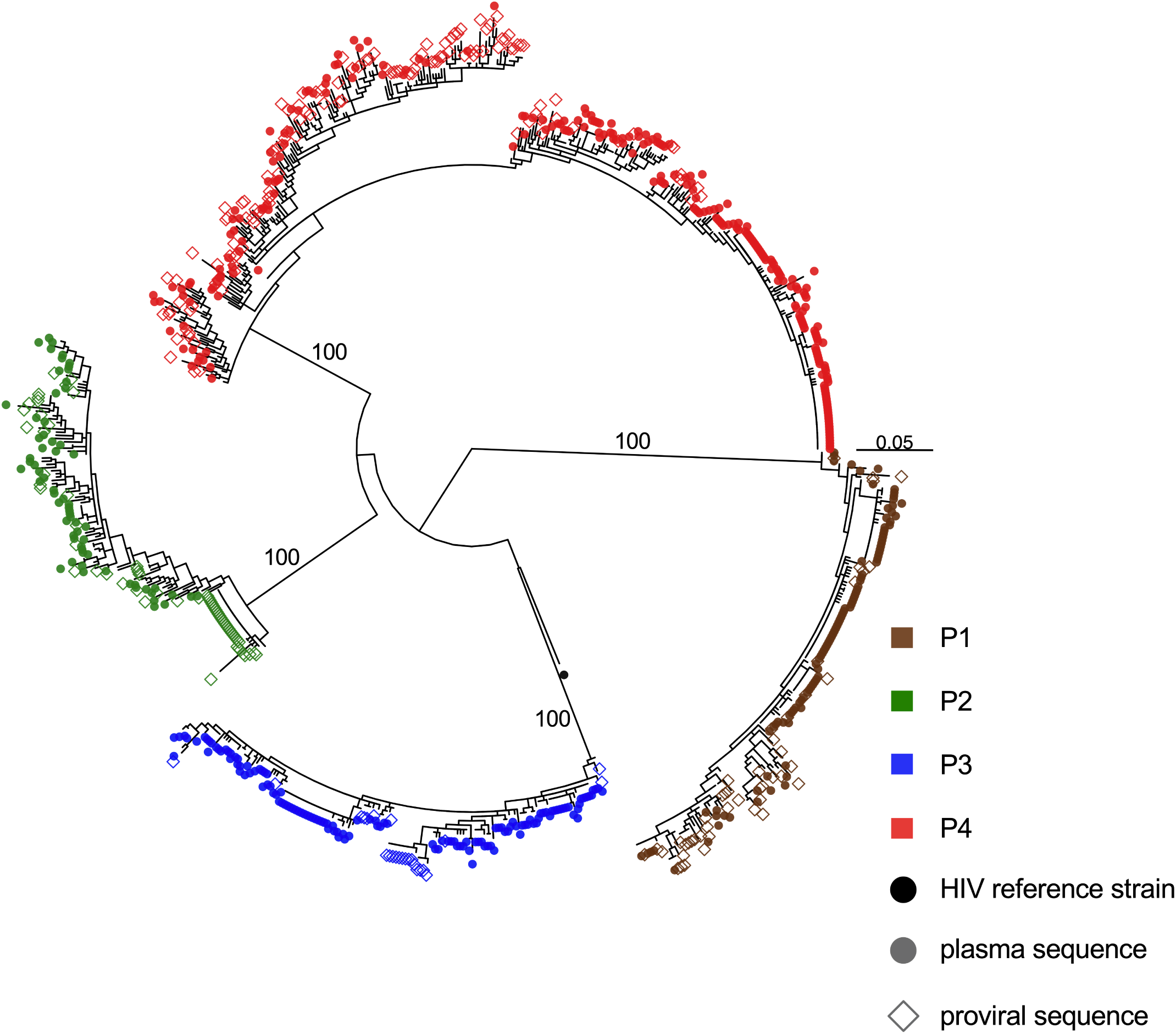
Between-host HIV phylogeny. Maximum likelihood phylogeny inferred from 546 plasma HIV RNA sequences (circles) and 267 proviral DNA sequences (open diamonds) isolated from participants. Numbers on internal branches indicate bootstrap values supporting within-host monophyletic clades. Scale in estimated substitutions per nucleotide site. The phylogeny is midpoint-rooted. The black dot represents the HIV-1 subtype B reference strain HXB2.

### Within-host HIV evolutionary reconstruction and proviral dating

We next characterized the diversity of proviruses persisting on ART, and inferred their ages phylogenetically (20). For each participant, we inferred a maximum likelihood phylogeny relating their plasma HIV RNA and proviral sequences, where the root represented the inferred transmitted founder event. We then fit a linear model relating the root-to tip distances of distinct pre-ART plasma HIV RNA sequences to their collection dates, where the slope of this line represents the within-host HIV *nef* evolutionary rate under a strict molecular clock assumption, and the x-intercept represents the phylogenetically-estimated infection date. We observed statistically significant within-host HIV evolution in plasma in all participants pre-ART, as defined by increasing divergence from the root, where the 95% confidence interval (CI) of the phylogenetically-estimated infection date captured the participant’s clinically estimated one in all cases (Table 2). We next describe each participant’s results in detail.

**Table 2:**
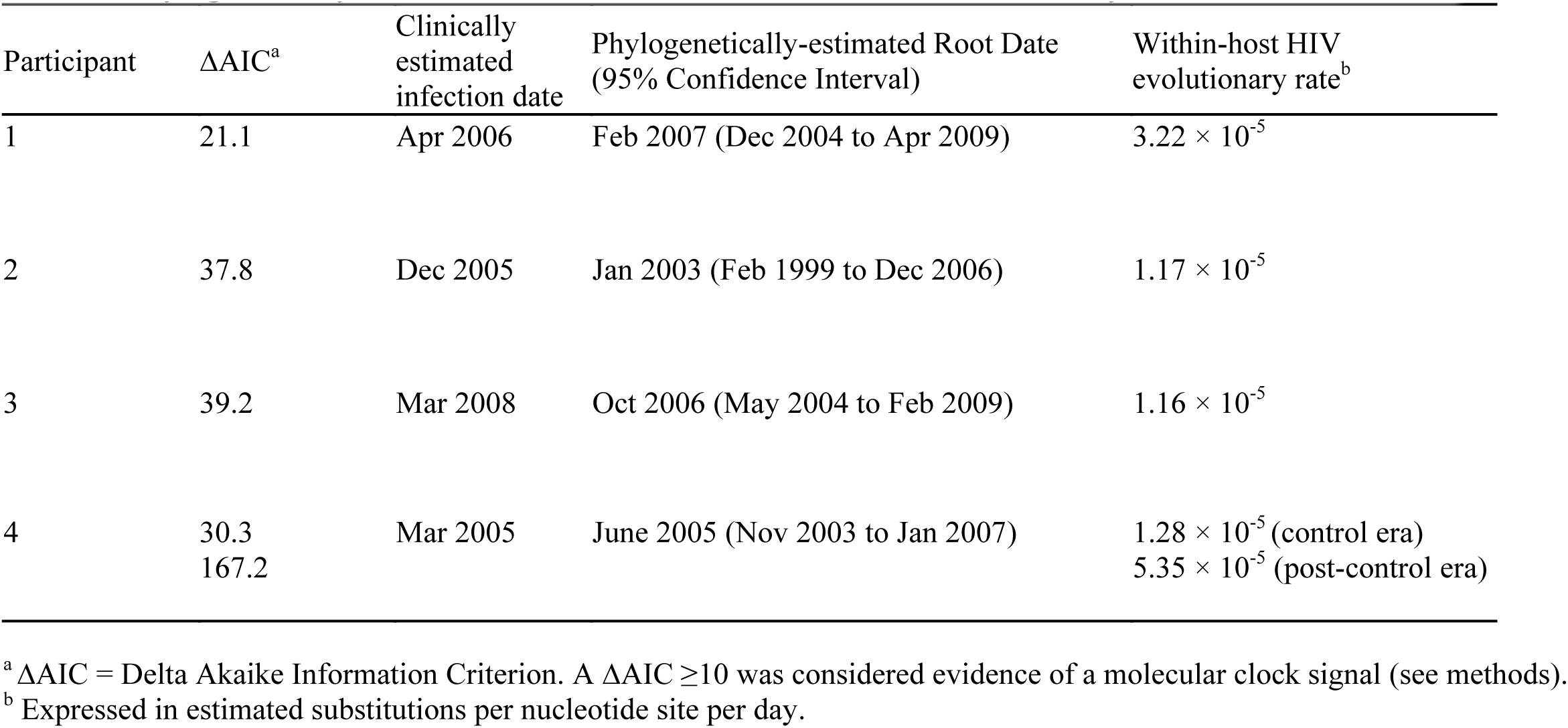
Phylogenetically-inferred root date estimates and within-host evolutionary rates

| Participant | $\Delta\text{AIC}^a$ | Clinically estimated infection date | Phylogenetically-estimated Root Date (95% Confidence Interval) | Within-host HIV evolutionary rate<sup>b</sup> |
| --- | --- | --- | --- | --- |
| 1 | 21.1 | Apr 2006 | Feb 2007 (Dec 2004 to Apr 2009) | $3.22 \times 10^{-5}$ |
| 2 | 37.8 | Dec 2005 | Jan 2003 (Feb 1999 to Dec 2006) | $1.17 \times 10^{-5}$ |
| 3 | 39.2 | Mar 2008 | Oct 2006 (May 2004 to Feb 2009) | $1.16 \times 10^{-5}$ |
| 4 | 30.3<br>167.2 | Mar 2005 | June 2005 (Nov 2003 to Jan 2007) | $1.28 \times 10^{-5}$ (control era)<br>$5.35 \times 10^{-5}$ (post-control era) |

<sup>a</sup> $\Delta\text{AIC}$ = Delta Akaike Information Criterion. A $\Delta\text{AIC} \geq 10$ was considered evidence of a molecular clock signal (see methods).
<sup>b</sup> Expressed in estimated substitutions per nucleotide site per day.

Participant 1 (EDI April 2006) maintained viremic control except for two plasma viral load measurements of 8075 and 2685 copies/ml in February and October 2008 respectively, prior to initiating ART in August 2010 (Figure 3A). Their proviral load on suppressive ART was 215 HIV copies/10^6^ CD4+ T cells, with 51% genetically intact (Figure 1; Table 1). We isolated 104 intact HIV RNA *nef* sequences from 8 pre-ART plasma timepoints spanning 2.4 years, of which 52 (50%) were distinct, and 48 proviral *nef* sequences sampled approximately two years post-ART, of which 40 (83%) were distinct (Table 1). The recovery of identical proviral sequences is consistent with clonal expansion of CD4+ T cells harboring integrated HIV (32–34), though we cannot definitively identify these as clonal since we only performed subgenomic HIV sequencing. Within-host phylogenetic analysis revealed increasing plasma HIV RNA divergence from the inferred root over time, along with non-synonymous substitutions that typify within-host HIV evolution (Figure 3B). Proviruses sampled on ART interspersed throughout the phylogeny except within the clade closest to the root, which comprised the earliest sampled plasma HIV RNA sequences. The linear model relating the root-to-tip distances of distinct pre-ART plasma HIV sequences to their collection dates yielded a pre-ART *nef* evolutionary rate of 3.22 × 10^-5^ substitutions per nucleotide site per day (Figure 3C; Table 2). Based on this, the oldest sampled provirus was estimated to have integrated in April 2008, 4 years prior to sampling (Figure 3D). Though the 95% CIs around the proviral date estimates are wide, these data nevertheless suggest that proviruses seeded throughout infection persisted during ART.

**Fig. 3:**
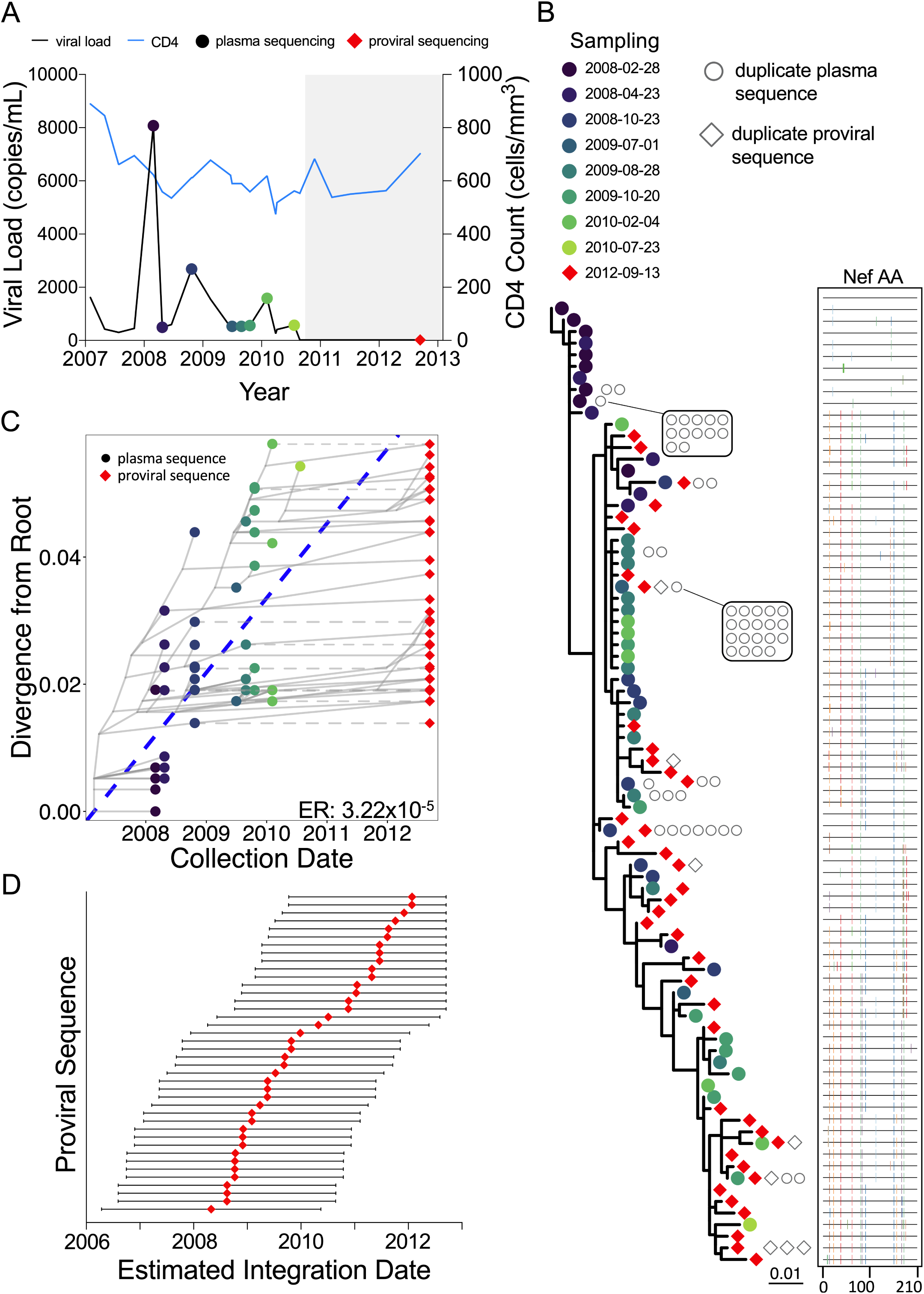
Participant 1. (A) Clinical history and sampling timeline. Throughout all figures, circles denote plasma HIV RNA sampling and diamonds denote HIV DNA sampling. Shading represents ART. (B) Maximum-likelihood within-host phylogeny and corresponding amino acid highlighter plot. In the phylogeny, colored circles denote distinct pre-ART plasma HIV RNA sequences and red diamonds indicate distinct proviral sequences sampled during suppressive ART. Sequences that were recovered repeatedly are shown as open grey circles (for plasma HIV RNA) and diamonds (for proviruses) adjacent to the relevant tip. The root represents the inferred most recent common ancestor of the dataset, representing the phylogenetically-inferred transmitted founder virus event. The highlighter plot is ordered according to the phylogeny and depicts *amino acid* sequences. The top sequence serves as the reference, where colored ticks in sequences beneath it denote non-synonymous substitutions with respect to the reference. (C) HIV sequence divergence-versus-time plot. The blue dashed line represents the linear model relating the root-to-tip distances of distinct pre-ART plasma HIV RNA sequences (colored circles) to their sampling times. This model is then used to convert the root-to-tip distances of distinct proviral sequences sampled during ART (red diamonds) to their original integration dates. The slope of the regression line, which represents the inferred within-host evolutionary rate (ER) in estimated substitutions per nucleotide site per day, is shown at the bottom right. Faint grey lines denote the ancestral relationships between HIV sequences. (D) Integration date point estimates (and 95% confidence intervals) for distinct proviral sequences recovered from this participant.

Participant 2 (EDI December 2005) maintained pVL <2000 copies/ml except for two measurements of 2160 and 3230 copies/ml in June 2010 and September 2013 respectively, and initiated ART in May 2014 (Figure 4A). Their total proviral load on ART was 580 HIV copies/10^6^ CD4+ T cells, with only 10% intact, the smallest intact proportion of all participants (Figure 1; Table 1). We recovered 67 HIV RNA *nef* sequences from 9 pre-ART plasma timepoints spanning more than four years, of which 60 (90%) were distinct, and 66 intact proviral *nef* sequences after 1.2 years on ART, of which 36 (55%) were distinct. The sequences closest to the root of this participant’s phylogeny were proviruses sampled on ART, consistent with these having integrated shortly after transmission (Figure 4B). Proviruses also interspersed throughout almost all descendant clades. Based on this individual’s calculated pre-ART *nef* evolutionary rate of 1.17 × 10^-5^ substitutions per nucleotide site per day (Figure 4C and Table 2), proviruses persisting on ART were inferred to have integrated on dates that spanned the individual’s entire pre-ART history (Figure 4D). One particular proviral sequence, recovered 26 times, was estimated to have integrated 6 years prior to sampling.

**Fig. 4:**
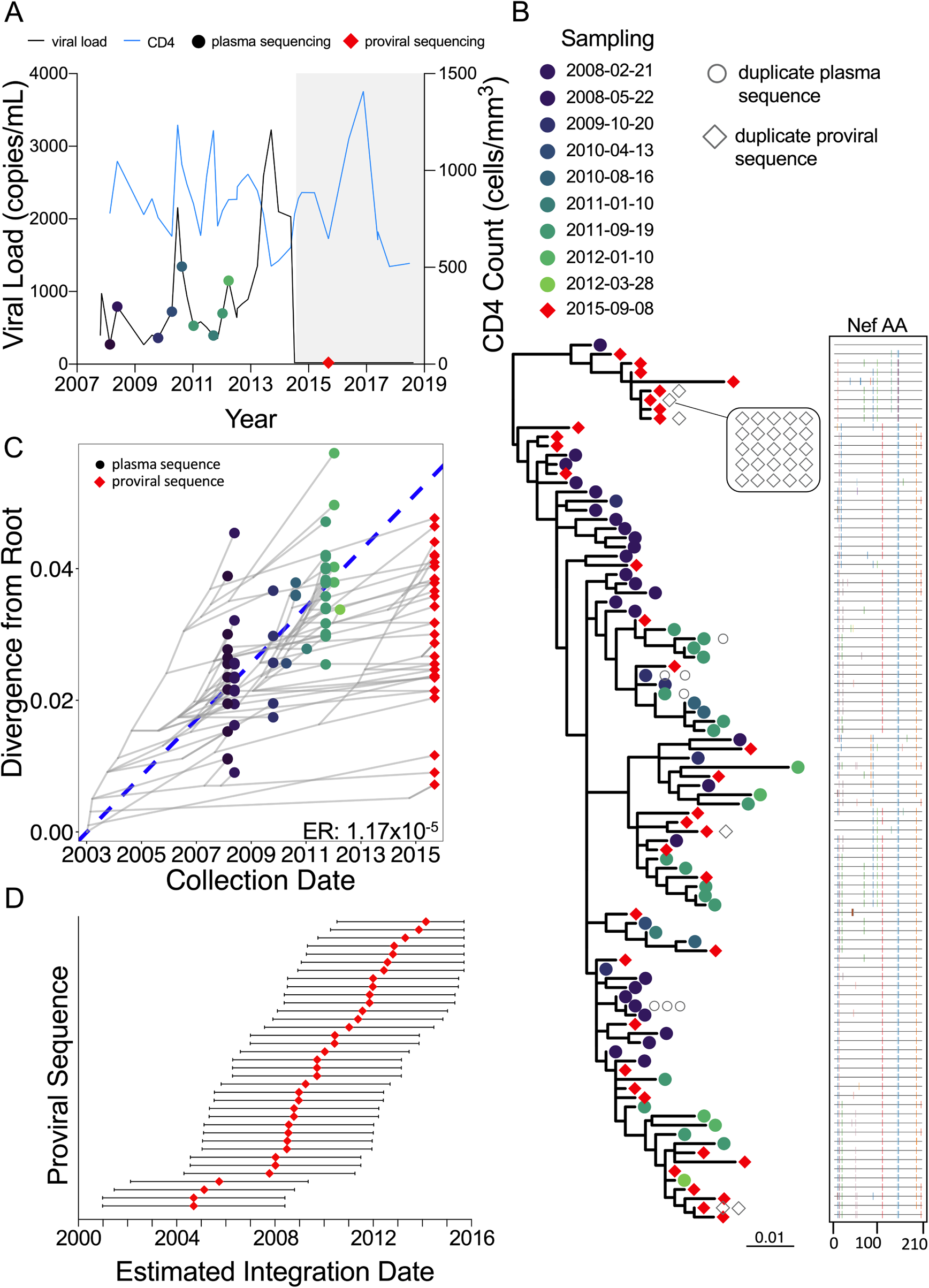
Participant 2. The panels are as described in the legend of Figure 3.

Participant 3 (EDI Mar 2008) continuously maintained pVL <1000 copies/ml before initiating ART in January 2012. Thereafter, pVL remained suppressed except during a treatment interruption when a pVL of 1610 copies/ml was recorded (November 2013), but unfortunately no plasma was available from that event from which to isolate HIV sequences. This participant had the smallest proviral burden of all (44 HIV copies/10^6^ CD4+ T cells), with an estimated 31% genetically intact (Figure 1; Table 1). We recovered 130 pre-ART plasma HIV *nef* sequences (55% distinct) from 7 pre-ART timepoints, where early plasma timepoints more frequently yielded identical HIV sequences, consistent with limited viral diversity following infection. We recovered only 24 proviral sequences (50% distinct) on ART, consistent with the small proviral burden. Despite maintenance of pVL<1000 pre-ART, the within-host phylogeny displayed significant molecular clock signal (1.16 × 10^-5^ substitutions per nucleotide site per day) where, similar to participant 2, proviruses sampled on ART interspersed throughout the tree and represented those closest to the root (Figures 5B, 5C). The earliest proviruses dated to around transmission while the latest dated to the viremic episode that occurred during the treatment interruption; the narrower 95% confidence intervals around these point estimates allow us to say with more certainty that this individual’s proviral pool spans a wide age range (Figure 5D). One sequence, recovered 9 times on ART and representing 38% of recovered proviruses in this individual, dated to around ART initiation.

**Fig. 5:**
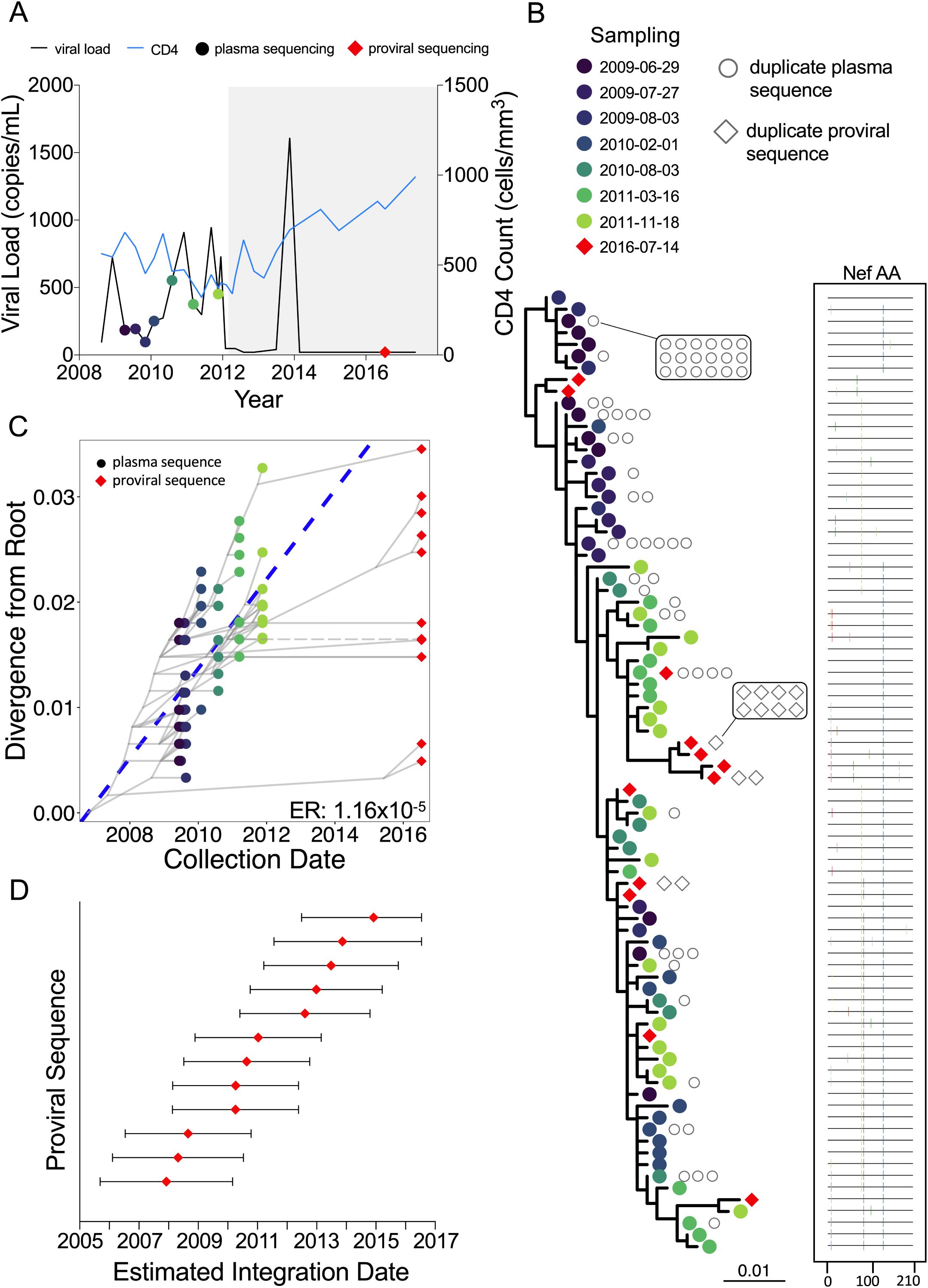
Participant 3. The panels are as described in the legend of Figure 3.

Participant 4 (EDI March 2005) initially controlled pVL to <2000 copies/ml for 3 years, but afterwards lost control, where pVL reached a maximum of 65,500 copies/ml before initiating ART in February 2013 (Figure 6A). Two linear models were calibrated, one each for the “control” and “post-control” periods, where the former was used to phylogenetically verify the clinically estimated infection date. We recovered 245 pre-ART plasma HIV *nef* sequences from 12 pre-ART timepoints (71% distinct), where, like participant 3, identical sequences were most frequently recovered from early timepoints (Figure 6B). We also recovered 129 proviral sequences during ART, a remarkable 92% of which were unique, indicating substantial proviral diversity in this individual. The linear model inferred from the viremic control period yielded strong molecular clock signal despite low viremia during this time, but no proviral sequences were recovered that interspersed with these very early sequences (Figures 6B, 6C). All recovered proviruses were therefore “dated” using the linear model representing the post-control period, yielding integration dates spanning this whole period (Figure 6C). Of note, the near-full-length genetically-intact provirus isolated during ART dated to 2011.

**Fig. 6:**
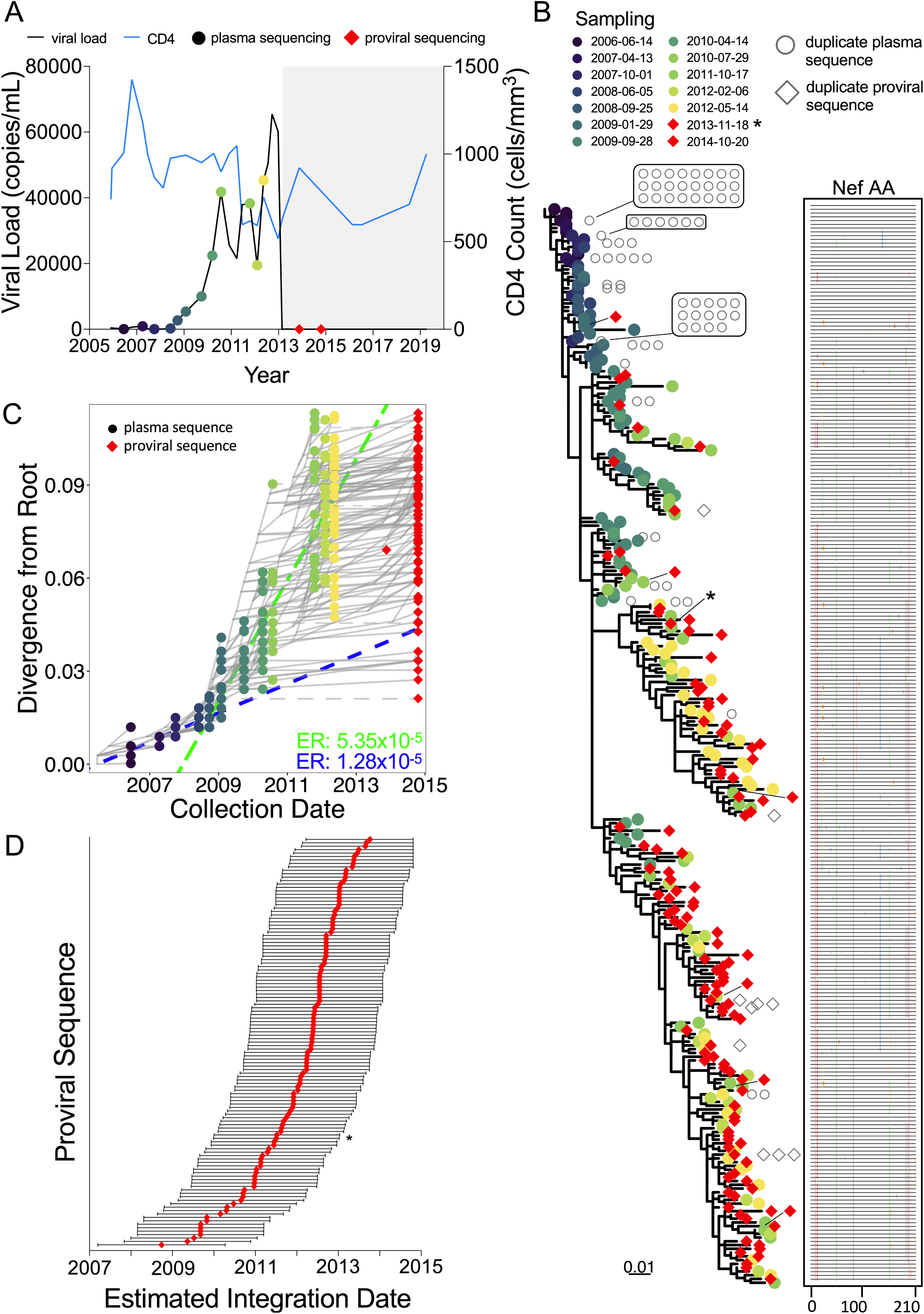
Participant 4. The panels are as described in the legend of Figure 3, with the following additions. Throughout the figure, the red diamond indicated by an asterisk represents the genomically-intact HIV sequence isolated from the sample remnants following the Intact Proviral DNA Assay (IPDA). In panel C, two linear models were fit to the data, encompassing the viremic control (blue dashed line) and the “loss of control” (green dashed line) periods. Corresponding evolutionary rates are shown in the bottom right corner in matching colours. The regression for the control period was performed using plasma timepoints 2006-06-14 to 2008-09-25, while post-control period regression was performed using plasma timepoints 2009-01-29 to 2012-05-14.

### Correlates of proviral burden and diversity during ART

Early ART limits HIV reservoir size (35–37), and positive correlations have been observed between the duration of uncontrolled viremia, and both HIV reservoir size and diversity in non-controllers (37–41). We therefore investigated the relationship between proviral burden, HIV diversity and cumulative viral load in our cohort. We observed a strong correlation between overall HIV genetic diversity in plasma pre-ART, and proviral diversity on-ART (Spearman’s ρ=1; p=0.08; Figure 7A). Furthermore, cumulative viral load, measured as log_10_ viremia copy-days during untreated infection, correlated strongly with both total proviral burden as measured by the IPDA and proviral diversity on ART (both Spearman’s ρ=1; p=0.08; Figures 7B, C).

**Fig. 7.**
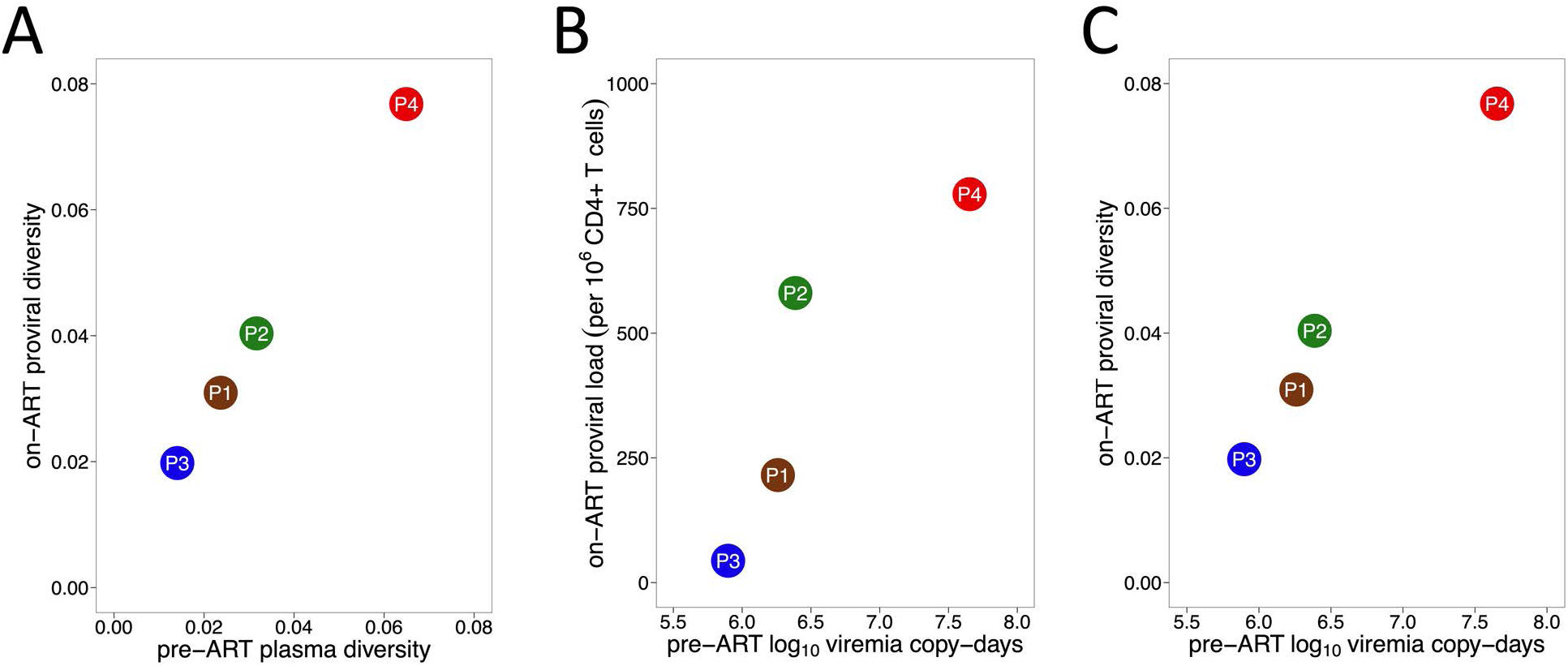
Correlates of proviral burden and diversity. (A) Relationship between pre-ART plasma HIV RNA diversity and proviral diversity on ART. (B) Relationship between total area under the plasma viral load curve pre-ART, measured in log_10_ viremia copy-days, and total proviral *burden* during ART. (C) Relationship between total area under the plasma viral load curve pre-ART and overall proviral *diversity* during ART, where the latter is measured as average patristic (tip-to-tip phylogenetic) distance between all distinct proviral sequences recovered from each participant.

### Inferring proviral turnover pre-ART: mathematical modeling approach

The decay rates (half-lives) of the proviral pool during untreated infection can be inferred from the age distributions of proviruses sampled on ART; this is because at the time of proviral sampling, the bulk of the proviral turnover had already occurred prior to therapy (19, 23, 24). We estimated pre-ART proviral half-lives in two ways: by adapting an existing mathematical model, and by estimating these rates directly from the data.

To do the former, we modified a dynamical mathematical model of HIV infection that describes within-host cell and virus concentrations over time within active and latent compartments (see methods and (23)). The model assumes that HIV sequences enter the reservoir at a rate proportional to their abundance in plasma, allowing us to model within-host proviral deposition in a personalized way. As our viral load data did not capture acute-phase dynamics, we merged each participants’ available data with acute phase dynamics estimated from the (limited) literature on HIV controllers (42–44) (see methods and **Table S1**) to model their proviral deposition. In a subsequent step, we then allowed these proviruses to decay at various constant, exponential rates up to the participants’ sampling date, yielding predictions of what the proviral age distribution would be, at time of sampling, for each decay rate tested.

The participants’ reconstructed viral load dynamics are shown in Figures 8A-D, while the model-predicted proviral compositions under different decay rates, depicted as the proportion of remaining proviruses that date to each year prior to ART (or in the case of participant 3, to the viremic episode that occurred during treatment interruption), are shown in Figures 8E-H. For participants 1-3, the model predicts that the bulk of the reservoir would have been deposited in the first year of infection (Figures 8E-G, yellow line); this is because the estimated acute-phase peak viral load is nearly 2 log_10_ higher than the subsequent setpoint. For participant 4, the model predicts that the bulk of the reservoir would have been deposited during the post-control period, because viral load was sustained at high levels during that time (Figure 8H, yellow line).

**Fig. 8.**
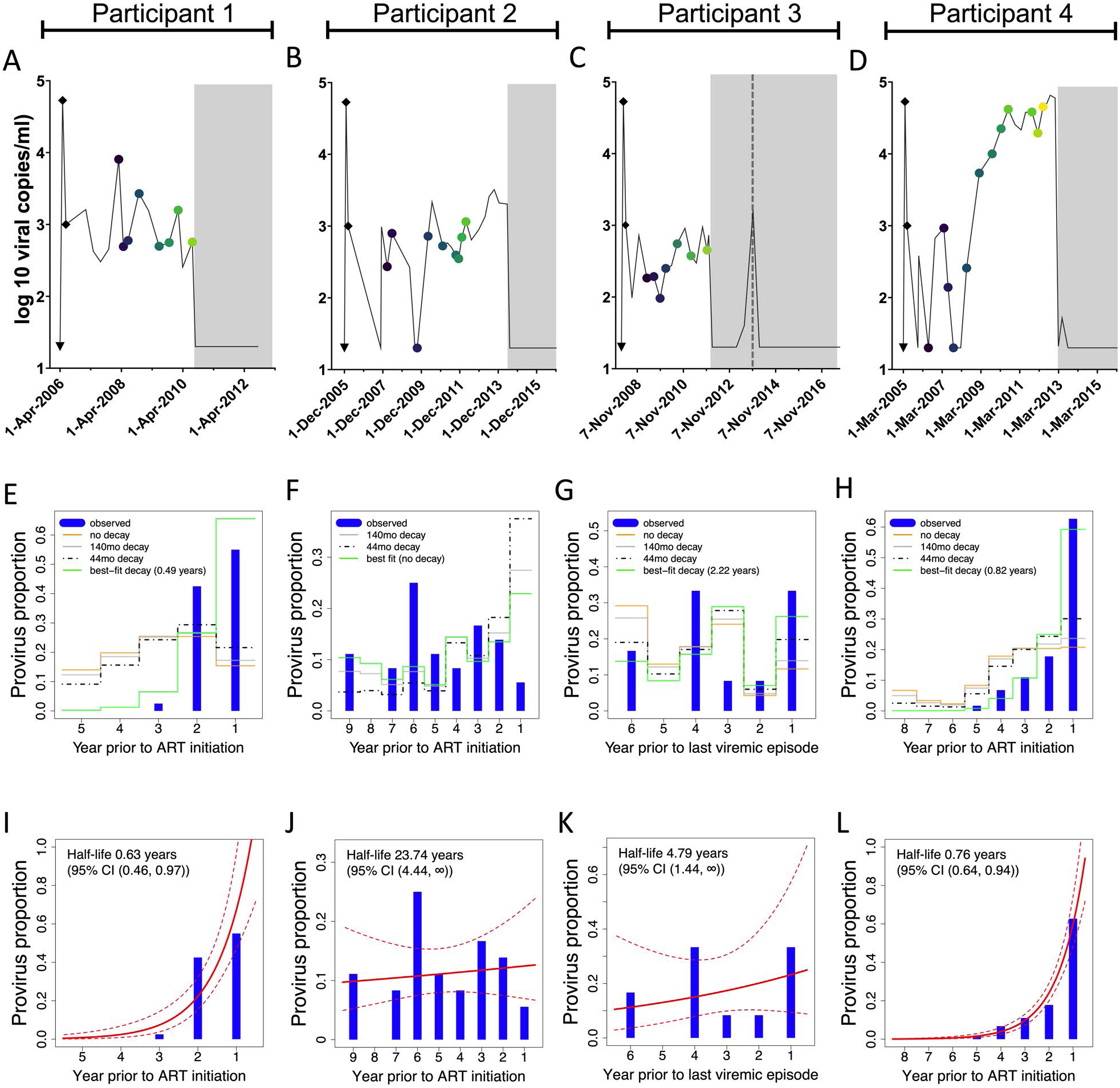
Model of reservoir composition. (A-D) Viral load histories, with engrafted “peak viremia” kinetics from the literature (shown as black diamonds), for participants 1 (panel A), 2 (panel B), 3 (panel C, where the dotted line represents the viremic episode after ART that serves as the reference point in this analysis) and 4 (panel D). (E-H) Phylogenetically-determined proviral ages, and predicted proviral age distributions under different rates of proviral decay, for participants 1-4. Blue histograms denote the proportion of each participant’s distinct proviruses that dated to each year prior to ART initiation, data that are derived from the integration date point estimates in Figures 3D, 4D, 5D and 6D, respectively. The yellow line indicates each participant’s model-predicted proviral deposition. The grey and dashed black lines predict what the proviral age distributions would be at the time of sampling, had proviral decay subsequently occurred under half-lives of 140 or 44 months following deposition (these half-lives represent published *on ART* estimates of total DNA (26) and replication-competent reservoir (25) decay, respectively). The green line represents the model-predicted proviral age distributions under the pre-ART proviral decay rate that best fit each participant’s observed data. (I-L) Best-fit half-lives estimated directly from each participant’s observed proviral age distributions using a Poisson generalized linear model (red line) along with 95% confidence intervals (dotted lines).

If we allow these deposited proviruses to decay at half-lives of 140 months (26) and 44 months (25), where these represent published estimates of DNA and replication-competent reservoir decay on ART respectively, the model predicts that, at time of sampling, participant proviral distributions would be skewed towards slightly younger ages (grey and black dotted lines). Notably, for participants 1, 3 and 4, these predicted proviral distributions fit their observed data (shown as the blue bars) quite poorly. As a last step, we identified the pre-ART proviral half-life that best fit each participants’ proviral distribution (green line). Participant 1’s was the shortest of all, only 0.41 (95% CI 0-1.03) years (Figure 8F). Estimated proviral half-lives for participants 3 and 4 were intermediate, while participant 2’s was nearly 9 years, with an upper 95% CI extending to 77 years.

While this model clearly illustrates how faster decay rates shift proviral distributions towards younger ages, it assumes that within-host cell and viral dynamics behave as parameterized by the model. The model-produced best-fit half-lives were also somewhat sensitive to imputed acute-phase viremia dynamics, with higher, longer peaks yielding the shortest half-lives (**Table S1**). The half-life estimates for participant 3, who controlled viremia the most successfully, were the most sensitive to these imputations: using a low, short peak for example yielded a half-life estimate of 3.04 (95% CI 0-18.08) years, nearly three times longer than that estimated in the primary analysis.

### Inferring proviral turnover pre-ART: direct approach

Recognizing the limitations of the dynamical model and our lack of capture of the participants’ actual acute-phase dynamics, we also estimated pre-ART proviral half-lives directly from observed proviral age distributions using a Poisson generalized linear model, an approach that does not incorporate clinical history information. The 95% CI around the half-life estimates produced by this method overlapped those produced by the dynamical model in all cases (Figures 8I-L). The point estimates for participants 1 and 2 were also very consistent, with the former again exhibiting the shortest proviral half-life of 0.63 (95% CI 0.46-0.97) years, and the latter exhibiting the longest of 23.74 (95% CI 4.44- ∞) years. Participant 4’s estimated pre-ART proviral half-life was 0.76 (95% CI 0.65-0.96) years, slightly shorter than that estimated with the dynamical model, while participant 3’s was 4.79 (95% CI 1.44- ∞) years, which was comparable to the dynamical model-derived estimate when using a low, short inferred acute-phase peak viral load (**Table S1**). Broadly however, the proviral pools that were skewed towards ART initiation produced short (<1 year) half-life estimates (participants 1 and 4) while those that dated more uniformly throughout infection produced substantially longer half-lives (participants 2 and 3).

### Sensitivity analyses: Alternate within-host phylogenies

We now present sensitivity analyses that test the robustness of our findings. In our primary analysis, we estimated proviral ages using a phylogeny inferred using the best-fit nucleotide substitution model. Each dataset however yielded models with similar predictive power (**Table S2**), so we re-computed proviral ages and pre-ART half-lives (using the Poisson generalized linear model) from a phylogeny inferred using each participant’s “next best fit” model. Results were broadly consistent with the primary findings: participants 1 and 4 again showed proviral distributions that were skewed to dates near ART, yielding pre-ART half-lives <1 year (**Figure S1)**. Participant 2’s alternative phylogeny dated a slightly larger proportion of proviruses to around ART initiation, yielding a shorter half-life than that estimated from the primary analysis. Nevertheless, both participant 2 and 3’s proviral distributions maintained an overall “flat” age distribution, yielding the longest estimated half-lives in the cohort. This suggests that our findings are not unduly influenced by the phylogenetic inference step.

### Inferring proviral age distributions and pre-ART half-lives without a molecular clock

In our primary analysis, we estimated proviral ages from root-to-tip phylogenetic distances by linear regression (20, 22). This assumes a strict molecular clock, which may not hold over long time periods (45), so we re-analyzed our data using two approaches that do not rely on this assumption. The first was “nearest neighbour” dating (21), where each provirus is assigned to the sampling date of the pre-ART plasma sequence closest to it in the tree (for consistency, the original tree was used). A limitation of this approach is that proviruses can only be assigned to dates when plasma HIV sequences were sampled. Nevertheless, the results were broadly comparable to the original ones (**Figure S2**, left side): participants 2 and 3 showed “flat” proviral age distributions and long estimated proviral half-lives, while participants 1 and 4 showed skewed proviral age distributions and proviral half-lives under one year.

We also used Least Squares Dating (LSD) (46), an approach that aims to minimize the variance between the tree branch lengths and sample dates, that has also recently been used to infer proviral sequence ages (47). We used the original rooted tree; here it may be useful to re-iterate that the root represents the location that maximizes the (Spearman’s) *correlation* between plasma HIV root-to-tip distances and sampling times, and thus does not rely on a strict clock. LSD also incorporates tree *topology* information (as opposed to just root-to-tip distances) and allows variable evolutionary rates over the tree’s edges. Overall, the LSD-inferred 95% CI were generally narrower and the within-host evolutionary rates slightly slower than those inferred using linear regression, but the estimated proviral ages and half-lives were again comparable to the original observations (**Figure S2,** right side). Taken together, these observations indicate that our findings are robust to a strict clock assumption.

### Accounting for identical sequences in proviral half-life estimates

When estimating rates of pre-ART proviral turnover we excluded identical proviral sequences under the assumption that these arose through clonal expansion rather than through independent integration events. Subgenomic HIV sequencing however cannot conclusively classify identical sequences as clonal across the whole HIV genome (48), so we re-analyzed our data using all sequences collected (Table 1). Results were again highly consistent with the original data (**Figure S3)**. Participant 1’s estimated proviral half-life when including 8 identical sequences was 0.61 [95% CI 0.46-0.9] years, which was very similar to the original 0.63 (95% CI 0.46-0.97) year estimate. Similarly, participant 2’s estimated half-life after including 30 identical sequences, the majority of which dated to a lineage that circulated 6 years prior to ART, was zero (*i.e.* no decay) which is comparable to the original 23.74 (95% CI 4.44-∞) year estimate. Our findings are therefore robust to the inclusion or exclusion of identical sequences.

## Discussion

Though modest in size, our longitudinal cohort of ART-treated viremic controllers allowed us to characterize proviral burden, diversity and dynamics in this understudied group. Our results indicate that the persisting proviral pools in these individuals can share considerable similarities in terms of size, diversity, and turnover with those of non-controllers on ART.

Within-host HIV evolutionary studies have revealed that most infections are initiated by a single transmitted founder virus that gives rise to increasingly diverse and divergent descendants (45, 49–54). The dynamics of within-host HIV evolution in controllers however remain somewhat unclear: while some studies performed on shorter timeframes reported increases in plasma HIV diversity and/or immune escape over time in elite (55–57) and viremic (58) controllers, others found limited or no evidence of evolution (12, 15, 59). Our first notable observation was that significant within-host HIV evolution occurred in all participants, as evidenced by ongoing divergence from a phylogenetically-inferred root, whose date was consistent with the clinically-estimated infection date in all cases. Also indicative of within-host evolution, non-synonymous substitutions consistent with escape from Human Leukocyte Antigen (HLA) class I-restricted cellular immune responses occurred even in participant 3, for whom pre-ART plasma viral loads never reached above 1000 HIV RNA copies/ml. In this participant who expresses HLA-A*03:01 and HLA-B*57:01, 25/25 sequences sampled in 2009 harbored the predicted HLA-B*57:01-restricted epitope AGNNAACAW at Nef codons 49-57 (60). By 2011, 8/27 sequences harbored A***<u>E</u>***NNAACAW which has a predicted >10-fold reduced binding affinity to B*57:01, a shift in frequency that was statistically significant (Fisher’s exact test; p=0.0044). Similarly, 25/25 sequences sampled in 2009 harbored the predicted A*03:01-restricted epitope KLVPVPEK at Nef codons 144-152, but by late 2011, 6/16 sequences harbored KLVPVPE***<u>E</u>***, which has a predicted >160-fold reduced binding affinity to A*03:01 (p=0.0069). Participants’ estimated within-host *nef* evolutionary rates, which ranged from 1.16 - 5.35 × 10^-5^ substitutions per nucleotide site per day assuming a strict clock, were also comparable to those reported in non-controllers using similar methods (20). Therefore, despite natural viremia control, significant within-host HIV evolution nevertheless occurs in this group in the absence of therapy.

Our results also revealed that total and genomically intact proviral DNA burdens in ART-treated viremic controllers can be substantial. While few studies have estimated reservoir size in controllers using the IPDA, a recent study reported 20-fold lower frequencies of total and intact proviral DNA in elite controllers (median of 30 and 1.6 copies/10^6^ CD4^+^ T cells respectively) compared to non-controllers on ART (median of 603 and 37 copies/10^6^ CD4^+^ T cells respectively) (61). The median total and intact proviral burdens in the present study were 398 and 84 copies/10^6^ CD4^+^ T cells respectively (or 215 and 58 copies/10^6^ CD4^+^ T cells respectively if we exclude participant 4 who lost viremic control), values that were more in line with those in non-controllers (30, 61, 62). Proviral loads in our study however were sampled only ∼1.5 years after ART on average. Longitudinal studies of treated viremic controllers on ART will therefore be required to confirm whether, similar to non-controllers, intact proviruses gradually decay on ART while total proviral DNA remains stable (30, 62, 63).

Our results also yield insights into on-ART proviral diversity and the length of time these sequences had persisted within-host. Consistent with a prior study that included viremic controllers (64), proviruses recovered in the present study were genetically diverse and frequently included archival lineages. Similar to studies of non-controllers (19–23), on-ART proviral age distributions varied: whereas participants 1 and 4’s proviral pools were skewed towards sequences that integrated in the years prior to ART, participants 2 and 3’s proviral pools spanned the infection course more evenly, and included proviruses that dated to shortly following infection. Despite these differences, the overall diversity of proviruses persisting on ART reflected the overall plasma HIV diversity generated prior to ART (Figure 7A). Moreover, the overall level of viral replication pre-ART correlated positively with both proviral burden and diversity on ART (Figures 7B-C). Our recovery of varying numbers of identical sequences also suggests that HIV controllers, like non-controllers, exhibit differential levels of clonal expansion (31–34, 65). Half of participant 3’s proviruses for example were identical to at least one other sequence, consistent with a report describing high identical proviral burdens in viremic controllers (64), while 83% of participant 1’s proviruses were distinct, a phenomenon that has also been reported in viremic controllers (58). Despite this heterogeneity, our observations confirm that diverse proviruses persist in those who control HIV prior to therapy, underscoring the importance of early ART even in this group.

Our study also extends our understanding of proviral turnover during untreated infection (19, 21, 23). While the half-life of the replication-competent and overall proviral pools on ART are on the order of ∼4 and >10 years respectively (25, 26, 30), recent data suggest that pre-ART proviral half-lives are shorter, with studies returning estimates of 8 (19, 24) and 25 months (23). Dynamic turnover during untreated infection helps explain why proviral pools are often enriched in sequences that integrated in the years immediately prior to ART (19, 21, 23), and has led to the hypothesis that ART dramatically slows this rate of turnover, thereby “stabilizing” the proviral pool in its immediate pretherapy state (21, 27). Indeed, participant 1 and 4’s skewed proviral distributions and short estimated pre-ART proviral half-lives resemble those of “typical” non-controllers (19, 23, 24), though participant 2 and 3’s flatter proviral age distributions and markedly longer estimated decay rates emphasize that proviral clearance is not rapid in all persons. While longer pre-ART half-lives have been observed in non-controllers (20), a recent study by our group identified an inverse relationship between set-point pVL and rate of pre-ART proviral turnover (24). Taken together with our observation that 2/4 participants exhibited relatively long pre-ART half-lives, this suggests that pre-ART proviral turnover may be slower in HIV controllers *on average*, though the data clearly underscore the heterogeneous nature of each individual’s persistent HIV pool.

Our study has some limitations. Our cohort is small due to the challenges of recruiting viremic controllers with known infection dates for longitudinal studies, and all participants were male. Though some aspects of the HIV reservoir likely differ between the sexes (66, 67), a recent study of non-controllers by our group revealed no significant differences in estimated pre-ART proviral half-lives between men and women, where intriguingly the participant with the “flattest” proviral distribution and longest pre-ART half-life was a female who nearly met the criteria for viremic control (24). This suggests that female controllers may not differ markedly from males in terms of proviral composition and pre-ART turnover, but additional studies are clearly needed to confirm this. As samples were limited and viral loads were low, near-full-genome HIV amplification was not feasible, nor was proviral integration site characterization, so we were not able to investigate genomic location and associated heterochromatin features recently characterized in elite controllers (8) in our cohort. Though *nef* is appropriate for within-host HIV evolutionary studies (20, 65, 68), sequencing of a subgenomic region does not allow us to classify recovered sequences as intact or defective, nor definitively classify identical sequences as clonally expanded. Our use of the IPDA to quantify intact (vs. defective) proviral burden mitigates this only partially. Our methods also assume that every provirus has equal probability of being sampled regardless of whether it is unique or a member of a clonally-expanded population (where we crudely estimate this probability to be <0.2%, as the blood collected represented ∼0.2% of average total body blood volume but subsequent DNA extraction and PCR efficiency is <100%). Sample availability also prevented us from investigating the mechanisms underlying participant 4’s loss of viremia control, though our observations that this individual’s (largely genetically intact) proviral pool dated exclusively to their post-control period suggests that this event triggered major increases in reservoir size and diversity, further underscoring the importance of timely ART, even in viremic controllers. Finally, the peak viremia kinetics that we drew from the literature (12, 42, 56) to use in the dynamical model may not have reflected actual *in vivo* kinetics, leading to uncertainty in our model-predicted proviral deposition and turnover rate estimates.

In conclusion, despite their natural ability to control HIV to low levels, significant within-host HIV evolution occurred in all participants pre-ART, giving rise to within-host proviral pools whose size and genetic diversity reflected this evolution. Pre-ART proviral dynamics were also heterogeneous: though two participants’ proviral pools were skewed towards recent integration dates consistent with rapid pre-ART turnover (19, 23, 24). Those of two others spanned a wide age range, consistent with much slower pre-ART proviral turnover. HIV remission and cure strategies will need to overcome within-host proviral diversity, even in individuals who naturally control HIV prior to therapy.

## Methods

### Study participants

Four viremic controllers who naturally maintained plasma viral loads (pVL) <2000 HIV RNA copies/ml for at least three years, and for whom an infection date could be reliably estimated, were studied. All participants were male with a median age of 55 years. Participants 1, 2 and 3 broadly maintained pVL < 2000 pre-ART whereas participant 4 lost viremic control prior to ART. The estimated dates of infection (EDI) for participants 1, 2, and 4, were calculated as the midpoint between the last seronegative and first seropositive HIV test dates, while for participant 3 the EDI was determined as March 2008, 3 months prior to diagnosis, due to a documented exposure followed by seroconversion-like illness. Participant nadir CD4+ T-cell counts ranged from 320-516 cells/mm^3^, while CD4+ T cells counts at ART initiation ranged from 401-626 cells/mm^3^. HLA class I types were: A*30:01-A*32:01/B*13:02-B*40:02/C*06:02-C*15:02 (participant 1); A*24:02-A*32:01/B*08:01-B*14:01/C*07:02-C*08:02 (participant 2); A*03:01-A*03:01/B*14:02-B*57:01/C*06:02-C*08:02 (participant 3); and A*03:01-A*32:01/B*14:02-B*39:01/C*08:02-C*12:03 (participant 4). All participants provided written informed consent. This study was approved by the Massachusetts General Hospital Research Ethics Board, with additional approvals for sample and data analysis approved by the Simon Fraser University and Providence Health Care/University of British Columbia Research Ethics Boards.

### Single genome HIV RNA and proviral amplification

Phylogenetic inference of proviral ages requires the longitudinal isolation of HIV RNA sequences pre-ART, alongside proviral DNA sequences sampled on ART, ideally using single-genome approaches. Because plasma viral loads were low and biological material was limited (only 10 million PBMCs per timepoint) full-length HIV amplification was not feasible, so *nef* was amplified for the reasons described in the results. Total nucleic acids were extracted from plasma using the BioMerieux NucliSENS EasyMag system (BioMerieux, Marcy-l’Étoile, France), while genomic DNA was extracted from PBMCs using the Purelink^®^ Genomic DNA kit (Invitrogen). For pre-ART plasma samples destined for HIV RNA *nef* amplification, 1 mL of sample was extracted, eluted in 60 μL and subjected to DNase I digestion (New England Biolabs Ltd, cat #: M0303S) to minimize the risk of amplifying HIV DNA in these low pVL samples. For on-ART PBMC samples destined for proviral *nef* amplification, cells were split into aliquots of 5 million cells before extraction and elution into 60µl.

HIV *nef* was amplified using limiting-dilution nested RT-PCR (for HIV RNA) or nested PCR (for proviral DNA) using high fidelity enzymes and sequence-specific primers optimized for amplification of multiple HIV group M subtypes (65, 68). For HIV RNA extracts, cDNA was generated using NxtScript Reverse Transcriptase (Roche). Next, cDNA as well as genomic DNA extracts were endpoint diluted such that ∼25-30% of the resulting nested PCR reactions, performed using the Expand^TM^ High Fidelity PCR system (Roche), would yield an amplicon. Primers (5’ -> 3’) used for cDNA generation/1^st^ round PCR were Nef8683F_pan (Forward; TAGCAGTAGCTGRGKGRACAGATAG) and Nef9536R_pan (Reverse; TACAGGCAAAAAGCAGCTGCTTATATGYAG). Primers used for 2^nd^ round PCR were Nef8746F_pan (Forward; TCCACATACCTASAAGAATMAGACARG) and Nef9474R_pan (Reverse; CAGGCCACRCCTCCCTGGAAASKCCC). Negative amplification controls were included in every run, and we also confirmed that plasma HIV RNA amplification did not occur in the absence of reverse transcription, indicating that DNAse treatment was effective. Amplicons were sequenced on an ABI 3730xl automated DNA analyzer. Chromatograms were basecalled using Sequencher v5.0 (Gene Codes) or the custom software RECall (69). *Nef* sequences that contained nucleotide mixtures, hypermutations (identified using Hypermut 2.0 (70)), suspected within-host recombinant sequences (identified using rdp4 Beta 95 (71)) or other defects were excluded from phylogenetic analysis, as were duplicate sequences (instead, the latter were added back at the data visualization stage). Within-host plasma and proviral *nef* sequences were codon-aligned using MAFFT v7 (72) implemented in HIV Align (https://www.hiv.lanl.gov/content/sequence/VIRALIGN/viralign.html) and manually edited in AliView v1.18 (73). Maximum likelihood phylogenies were inferred from aligned, gap-stripped within-host sequence datasets using W-IQ-TREE (74) following automated model selection with ModelFinder (75) using an Akaike Information Criterion (AIC) selection criterion (76). Best-fit models are reported in **Table S2**. Our primary analysis used a phylogeny inferred using each participant’s top best-fit model, but a sensitivity analysis was performed using each participant’s next best fit model (**Figure S1**). Patristic (tip-to-tip phylogenetic) distances were extracted using the cophenetic.phylo function from the R package ape (v5.3) (77).

### HIV proviral full genome amplification and sequencing

Single-template, near-full-length proviral amplification was performed on DNA extracted from CD4+ T cells by nested PCR using Platinum Taq DNA Polymerase High Fidelity (Invitrogen) such that ∼25% of the resulting PCR reactions yielded an amplicon. First round primers were: Forward - AAATCTCTAGCAGTGGCGCCCGAACAG, Reverse - TGAGGGATCTCTAGTTACCAGAGTC. Second round primers were: Forward - GCGCCCGAACAGGGACYTGAAARCGAAAG, Reverse- GCACTCAAGGCAAGCTTTATTGAGGCTTA. Reactions were cycled as follows: 92°C for 2 minutes; 10 cycles of (92°C for 10 seconds, 60°C for 30 seconds and 68°C for 10 minutes); 20 cycles of (92°C for 10 seconds, 55°C for 30 seconds and 68°C for 10 minutes); 68°C for 10 minutes (78, 79). Amplicons were sequenced using Illumina MiSeq technology and *de novo* assembled using the custom software MiCall (https://github.com/cfe-lab/MiCall) which features an in-house modification of the Iterative Virus Assembler (IVA) (80).

### Within-host phylogenetic inference and proviral age reconstruction

We used a published within-host phylogenetic approach to infer proviral sequence ages (20). Briefly, for each participant we inferred a maximum likelihood phylogeny relating within-host longitudinal pre-ART plasma HIV RNA and proviral sequences sampled on ART. We then exhaustively re-root each tree to identify the root location that maximizes the (Spearman’s) correlation between the root-to-tip distances and collection dates of the pre-ART plasma HIV RNA sequences. Here, the root represents the inferred most recent common ancestor (MRCA) of the within-host dataset (*i.e.* the inferred transmitted founder virus). We then fit a linear model relating the collection dates of the plasma HIV RNA sequences to their divergence from the root, where the slope represents the average host-specific rate of HIV evolution prior to ART and the x-intercept represents the phylogenetically-estimated MRCA date. For participant 4, two regression lines were fit: one each for their “viremic control” and “post-control” eras.

The integration dates of the proviral sequences sampled during ART, along with their 95% confidence intervals (CI), are then estimated from their divergence from the root using the linear regression. We assessed molecular clock and model fit using a delta (Δ) Akaike Information Criterion (ΔAIC) (76), computed as the difference between the AIC of the null model (no evidence of within-host evolution, *i.e.* a zero slope) and the AIC of the linear regression. A within-host phylogeny was deemed to have a sufficient molecular clock signal if the linear regression produced a ΔAIC≥10, (which corresponds to a p-value of 0.00053 when using a log-likelihood ratio test), and the 95% confidence interval of the estimated root date contained or preceded the first sampling point. The proviral dating framework is available as a web-service, at https://bblab-hivresearchtools.ca/django/tools/phylodating, with open source code available at https://www.github.com/cfe-lab/phylodating. As described in the results, sensitivity analyses were also performed where proviruses were phylogenetically “dated” using two published methods that do not rely on a strict molecular clock assumption (**Figure S2**).

### Intact Proviral DNA Assay (IPDA)

Each participant’s first PBMC sample on ART was used to estimate total and intact proviral DNA burdens using the Intact Proviral DNA Assay (IPDA) (29). As this method reports proviral burdens in terms of HIV copies per million CD4+ T cells, we first isolated CD4+ T cells from 10M PBMC by negative selection using the EasySep Human CD4+ T cell Enrichment Kit (STEMCELL Technologies, Cat #: 19052). This yielded a mean of 1.6 (range 1.3-2.0) million CD4+ T cells per participant, which were then extracted using the QIAamp DNA Mini Kit (Qiagen) with precautions to minimize shearing. In the IPDA, HIV and human DNA quantification reactions (where the latter target the human RPP30 gene) are conducted in parallel, where copies are normalized to the quantity of input DNA and subsequently corrected for DNA shearing. In each ddPCR reaction, 7ng (RPP30) or a median 588ng (IQR 488-697ng) (HIV) of genomic DNA was combined with ddPCR Supermix for Probes (no dUTPs, BioRad), primers (final concentration 900nM, Integrated DNA Technologies), probe(s) (final concentration 250nM, ThermoFisher Scientific), XhoI restriction enzyme (New England Biolabs) and nuclease free water. Human RPP30 primer and probe sequences (5’ -> 3’) are: RPP30 Forward Primer- GATTTGGACCTGCGAGCG, RPP30 Probe- VIC- CTGACCTGAAGGCTCT- MGBNFQ, RPP30 Reverse Primer- GCGGCTGTCTCCACAAGT; RPP30 Shear Forward Primer- CCATTTGCTGCTCCTTGGG, RPP30 Shear Probe- FAM- AAGGAGCAAGGTTCTATTGTAG- MGBNFQ, RPP30 Shear Reverse Primer- CATGCAAAGGAGGAAGCCG. The default (29) HIV primers and probes were: Ψ Forward Primer- CAGGACTCGGCTTGCTGAAG, Ψ Probe- FAM- TTTTGGCGTACTCACCAGT- MGBNFQ, Ψ Reverse Primer- GCACCCATCTCTCTCCTTCTAGC; *env* Forward Primer- AGTGGTGCAGAGAGAAAAAAGAGC, *env* Probe- VIC-CCTTGGGTTCTTGGGA- MGBNFQ, anti-Hypermutant *env* Probe- CCTTAGGTTCTTAGGAGC- MGBNFQ, *env* Reverse Primer- GTCTGGCCTGTACCGTCAGC.

The published HIV primers/probes however can sometimes fail to detect autologous HIV sequences due to naturally-occurring polymorphism (81, 82), thus requiring autologous primer/probes. For this reason, targeted HIV sequencing of the IPDA Ψ and *env* regions was performed for all participants prior to IPDA measurement, and autologous primers/probes substituted where necessary. The substituted primer and probe sequences (5’ -> 3’) were: Participant 2 Ψ Probe- FAM- TTTCAGCGTACTCACCAGT, Participant 3 Ψ Probe- FAM- ATATGGCGTACTCACCGGT, Participant 1 Ψ Probe- FAM- AATTGGCGTACTCACCAGT, Participant 1 Ψ Probe- FAM- AATTGGCGTACTCACCAGC, Participant 1 *env* Forward Primer- GGTGGTGCAGAGAGAAAAAAGAGC, Participant 1 *env* Reverse Primer- GCTGACGGCACAGGCCAGGC, Participant 3 *env* Reverse Primer-GCTGACGGTACAGGCCAGAT. Droplets were prepared using the Automated Droplet Generator and cycled as previously described (29). Droplets were analyzed on a QX200 Droplet Reader (BioRad) using QuantaSoft software (BioRad, version 1.7.4), where the results of a minimum of 8 technical replicates, comprising a median of 589,820 (IQR 501,539-721,871) cells assayed in total, were merged prior to analysis. Intact HIV-1 copies (Ψ and *env* double-positive droplets) were corrected for DNA shearing based on the frequency of RPP30 and RPP30-Shear double positive droplets. The median DNA shearing index, which measures the proportion of sheared DNA in a sample, was 0.40 (IQR 0.39-0.43), which is in line with acceptable levels in this assay (29, 81).

### Inference of pre-ART proviral half-lives: dynamical mathematical model

In order to infer pre-ART proviral half-lives, each participant’s sampled proviruses were grouped into “bins” by their estimated year of integration (one “bin” per year preceding ART initiation). For simplicity, proviruses whose integration date point estimate fell after the ART initiation date (or in the case of participant 3 the date of the last viremic episode), were assigned to the ART initiation (or viremic episode) date (all proviruses in this category had 95% confidence intervals that overlapped this date). In the primary analysis, proviral half-life inference was only performed on distinct proviral sequences under the assumption that “replicate” sequences arose through clonal expansion and not through individual integration events, but a sensitivity analysis was performed including all proviral sequences (**Figure S3**).

We estimated host-specific pre-ART proviral half-lives two ways: using a dynamical mathematical model (23), and directly from the data. To do the former, we implemented a published mathematical model that describes each individual’s untreated HIV infection via a set of ordinary differential equations that model the dynamics of the concentrations of cells and virus in each individual over time (23). The model includes susceptible target cells, actively and latently infected cells that can produce viable virus, actively and latently infected cells that *cannot* produce viable virus, the virus itself, and an immune response. The model assumes that HIV sequences enter the latent proviral pool at a rate proportional to their abundance in plasma at the time. In a subsequent independent step, proviruses are then allowed to “decay” out of this pool at various exponential rates, to produce predictions of what the proviral age distribution would be at ART initiation if decay had occurred at this rate. We reproduce the model here.

Let *S* represent the susceptible compartment. Let *A*_*P*_ and *L*_*P*_ represent actively and latently infected cells that are productively infected and produce viable virus; likewise let *A*_*U*_ and *L*_*U*_ represent actively and latently infected cells that are *unproductively* infected and cannot produce viable virus. Let *V* represent viremia and let *E* represent the adaptive immune response. The following dynamical system describes within-host HIV infection kinetics:

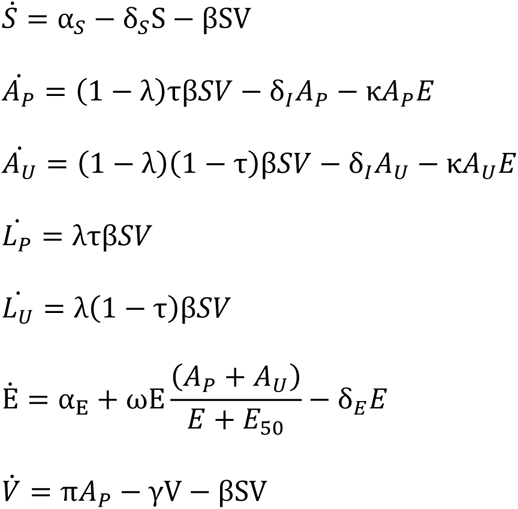

The parameters used were: susceptible cell creation rate *α*_*S*_ = 70 cells μL^−1^day^−1^; susceptible cell death rate δ_*S*_ = 0.2 *day*^−1^; viral infectivity β = 10^−4^ μL viral RNA copies^−1^day^−1^; probability of productively infectious virions τ = 0.05; probability of latency λ = 10^−4^; actively infected cell death rate δ_*I*_ = 0.8 day^−1^; viral burst size π = 50000 viral RNA copies cell^−1^day^−1^; viral clearance rate γ = 23 day^−1^; initial adaptive precursor frequency α_*E*_ = 10^−4^cells μL^−1^day^−1^; adaptive immune killing rate κ = 0.3 μL cells^−1^day^−1^; adaptive immune recruitment rate ω = 1.6 *day*^−1^; adaptive immune clearance rate δ_*E*_ = 0.002 day^−1^; adaptive immune 50% saturation constant (*i.e.* the number of infected cells required for a half-maximal cytolytic expansion rate (83)) *E*_50_ = 250 cells μL^−1^.

The following initial conditions were used as per the original publications (23, 83). The initial viral load value was *V*(0) = 0.03 viral RNA copies per μL, which is the detectable plasma viral load limit of a typical assay when converted from the conventional 30 HIV RNA copies/mL to viral RNA copies per microlitre. Also let *I*_0_ = *V*(0)γ/π = 1.38 × 10^−5^; this represents a quasistatic approximation for the infected cells.

- *S*(0) = α_*S*_/δ_*S*_ = 70/0.2 = 350
- *A*_*P*_(0) = *I*_0_τ(1 − λ) = 6.89931 × 10^−7^
- *A*_*U*_(0) = *I*_0_(1 − 𝜏)(1 − 𝜆) = 1.310869 × 10^−5^
- *L*_*P*_(0) = *I*_0_τλ = 6.9 × 10^−11^
- *L*_*U*_(0) = *I*_0_(1 − τ)λ = 1.311 × 10^−9^
- *E*(0) = α_*E*_/δ_*E*_ = 0.0001/0.002 = 0.05
- *V*(0) = 30/1000 = 0.03

Note that *S*(0) and *E*(0) are the equilibrium values for the system in the absence of virus.

As described above, reservoir creation is assumed to be proportional to plasma viral load over time, with the probability of a virus entering the latent pool (defined by λ above) remaining constant over time. Unfortunately, peak viremia was not captured in our participants’ measured viral load data. Therefore, to recreate each participant’s *in vivo* plasma viral dynamics as realistically as possible, we merged each participant’s available pVL data to acute phase dynamics observed in viremic controllers (42), and we also performed sensitivity analyses using peak viremia kinetics observed in elite controllers (43, 44) (**Table S1**). Using each participant’s reconstructed viral load history and the above equations, we modeled proviral deposition into their latent pool. We then grouped the resulting proviruses by their year of creation.

In a separate step, we then allowed each group of latent proviruses to decay exponentially, up until each participant’s proviral sampling date, under half-lives ranging from 30 to 6000 days, in increments of 30 days. This produced a series of 200 predicted proviral distributions per participant that represented what proportion of proviruses would still remain from each creation year, assuming decay at the stated rate. For context, we also applied decay rates of 44 and 140 months, which represent the half-life of the replication-competent reservoir (25) and total proviral pool (26) during suppressive ART, respectively. Note that the model assumes that, after ART, the overall size of the proviral pool decreases but the relative proportions of sequences in each group remains the same; as such, model predictions would be the same regardless of when the proviral pool is sampled on ART. Finally, we identified the best-fit proviral decay rate to each participant’s *observed* proviral age distribution by maximum likelihood, and used standard theory to identify 95% confidence bounds.

### Inference of pre-ART proviral half-lives: Poisson Linear Model

To estimate the reservoir half-life directly from the data, we again grouped each participant’s observed proviral sequences by their age, in years, relative to ART initiation (or for participant 3, the date of their last viremic episode). We will refer to the bin containing proviruses from *t* − 1 to *t* years old as “bin *t*”, where we make the simplifying assumptions that all proviruses in bin *t* are exactly *t* years old. We then applied a Poisson generalized linear model with the canonical natural logarithm link function to the binned counts, using the age of the bin *t* as the predictor. This choice can be justified as follows: Let *t*_1/2_ be the proviral half-life. Assuming the participant’s proviral reservoir decays at an exponential rate, we would expect that the size of bin *t* would be approximately

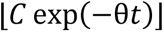

where *C* represents the initial size of the age bin as if there was no decay, and θ = ln(2) /*t*_1/2_.

Now, we assume that every provirus has the same small independent probability *p* of being sampled. If so, we would then expect the number of observed proviruses from bin *t* to be binomially distributed with ⌊*C* exp(−𝜃*t*)⌋ trials and success probability *p*. Assuming that *p* is small, we can approximate the distribution with a Poisson distribution with parameter

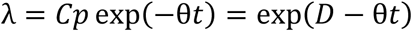

where 𝐷 = ln(*Cp*), or in other words,

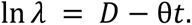

which is precisely the setup required for a Poisson generalized linear model with natural logarithm link function.

### Statistical Analyses

Spearman’s correlation was used to assess the relationship between continuous parameters. Statistical analysis was performed in Prism (version 9) software.

## Data Availability

GenBank accession numbers for sequences reported in this study are MW781931 to MW782197 (proviral DNA) and MW782198 to MW782743 (plasma HIV RNA).

## Acknowledgements

The authors gratefully thank the study participants without whom this research would not be possible. The authors also thank Dr. Daniel Reeves, the creator of the dynamical mathematical model, for helpful correspondence. We are also grateful to Daniel Worrall for his assistance with clinical data access.

## Funding Statement

This work was supported in part by the Canadian Institutes of Health Research (CIHR) through a project grant (PJT-159625 to ZLB and JBJ) and a focused team grant (HB1-164063 to ZLB and MAB). This work was supported in part by the National Institutes of Health (NIH) under award number NIHA127029 (to ZLB) and the Martin Delaney ‘BELIEVE’ Collaboratory (NIH grant 1UM1AI26617 to ZLB and MAB), which is supported by the following NIH Co-Funding and Participating Institutes and Centers: NIAID, NCI, NICHD, NHLBI, NIDA, NIMH, NIA, FIC, and OAR. Research reported in this publication was supported by the National Institute of Allergy and Infectious Diseases of the National Institutes of Health under Award Number UM1AI164565. Research reported in this publication was also supported in part by the National Institute Of Allergy And Infectious Diseases of the National Institutes of Health under Award Number UM1AI164565 (Martin Delaney ’REACH’ collaboratory; to ZLB and MAB). The content is solely the responsibility of the authors and does not necessarily represent the official views of the National Institutes of Health. FHO is supported by a Ph.D. fellowship from the Sub-Saharan African Network for TB/HIV Research Excellence (SANTHE), a DELTAS Africa Initiative [grant # DEL-15-006]. The DELTAS Africa Initiative is an independent funding scheme of the African Academy of Sciences (AAS)’s Alliance for Accelerating Excellence in Science in Africa (AESA) and supported by the New Partnership for Africa’s Development Planning and Coordinating Agency (NEPAD Agency) with funding from the Wellcome Trust [grant # 107752/Z/15/Z] and the UK government. The views expressed in this publication are those of the authors and not necessarily those of AAS, NEPAD Agency, Wellcome Trust or the UK government. AS and BRJ are supported by CIHR Doctoral Research Awards. NNK is supported by a CIHR Vanier Doctoral Award. RLM is supported by a CIHR MSc award. MAB holds a Canada Research Chair, Tier 2 in Viral Pathogenesis and Immunity. ZLB is supported by a Scholar Award from the Michael Smith Foundation for Health Research.

## Supplemental Files

**Table S1: Within-host proviral half-life estimates from primary and sensitivity analyses.**

**Table S2**: **Best-fit nucleotide substitution models.**

**Fig. S1 Inferring proviral age distributions and pre-ART half-lives from alternate within-host phylogenies.** (A) Divergence vs. time plot for participant 1, inferred using a TVM+F+I+G4 nucleotide substitution model, which represents this participant’s next best fit model as shown in Table S2. As in the main manuscript figures, the blue dashed line represents the linear model relating the root-to-tip distances of distinct pre-ART plasma HIV RNA sequences (colored circles) to their sampling times; this line is used to convert the root-to-tip distances of distinct proviral sequences sampled during ART (red diamonds) to their integration dates. The slope of the line, which represents the within-host evolutionary rate (ER) in estimated substitutions per nucleotide site per day, is shown on the plot. Faint grey lines denote the ancestral relationships between HIV sequences. (B) Integration date point estimates and 95% confidence intervals for distinct proviral sequences recovered from participant 1, as inferred using the alternate phylogeny. (C) Proviral age distribution (blue histogram), and and associated best-fit half-life estimated using a Poisson generalized linear model (red line, with 95% CI shown as dotted line). (D-F) results from participant 2’s alternate phylogeny, inferred using a TPM2+F+I+G4 model. (G-I) results from participant 3’s alternate phylogeny, inferred using a TVM+F+I+G4 model. (J-L) results for participant 4’s alternate phylogeny, inferred using a GTR+F+I+G4 model. As in the primary analysis, two regression lines were fit, one each for the “control” (blue line and ER) and “post-control” eras (green line and ER), where all proviruses were dated using the second regression.

**Fig. S2 Inferring proviral age distributions and pre-ART half-lives without a molecular clock.** Left side: The first column shows the proviral integration date estimates inferred from each participant’s original phylogeny using the “nearest neighbor” dating method, which dates each provirus to the pre-ART plasma sample closest to it in the tree. This approach can only date proviruses to the specific dates on which plasma HIV RNA sequences were sampled, and therefore produces no 95% CI. The second column shows the proviral age distributions inferred using the nearest neighbor approach (blue histograms), and the associated best-fit half-life estimated using a Poisson generalized linear model (red line, with 95% CI shown as dotted line). Right side: The first column shows the divergence vs. time plots produced by Least Squares Dating (LSD). Here, all plasma HIV RNA sequences are shown in black with the evolutionary relationships between sequences shown as grey solid lines. A dotted horizontal line connects each distinct provirus (red diamond) to its position in the phylogeny (shown as a red cross), where its inferred date can be read on the x-axis. The inferred evolutionary rate, in estimated substitutions per nucleotide site per day, is shown at the bottom right of each plot. The second column shows the integration date point estimates and associated 95% confidence intervals for all distinct sampled proviruses. The third column shows the best-fit half-lives estimated directly from the LSD-inferred proviral age distributions.

**Fig. S3 Accounting for identical sequences in proviral half-life estimates**. Left column: Each participant’s original divergence vs. time plot is shown, with identical proviruses annotated outside the plot border as red diamonds, >5 duplicate sequences are indicated using numbers. Middle column: Proviral age distributions computed using distinct sequences only (blue histograms), and associated best-fit half-life estimated using a Poisson generalized linear model (red line, with 95% CI shown as dotted line). These are the same results as shown in Figures 8I-L. Right column: Proviral age distributions and associated best-fit half-lives when considering all proviral sequences.

## Notes

### Competing Interest Statement

The authors have declared no competing interest.

### Summary of Updates

Alternate within-host phylogenies: the primary analyses estimated proviral ages using a phylogeny inferred using the best-fit nucleotide substitution model. Each dataset however yielded models with similar predictive power. In the updated version we re-computed proviral ages and pre-ART half-lives (using the Poisson generalized linear model) from a phylogeny inferred using each participant's "next best fit" model yielding results that were consistent with primary findings. Secondly, we inferred proviral age distributions and pre-ART half-lives using two approaches that do not rely on the assumption of a molecular clock - namely, nearest neighbor "dating" method and least squares "dating" method. Results from this sensitivity analysis indicate that our findings were robust to a strict clock assumption. Lastly, in our primary analysis we excluded identical proviral sequences under the assumption that these arose through clonal expansion rather than through independent integration events. Subgenomic HIV sequencing however cannot conclusively classify identical sequences as clonal across the whole HIV genome, so we re-analyzed our data using all sequences collected yielding results that were consistent with the original data. The results from these sensitivity analyses are now presented as supplementary Figures S1 - S3.

