## Supplemental Table 1 for "HIV proviral burden, genetic diversity, and dynamics in viremic controllers who subsequently initiated suppressive antiretroviral therapy"

**Table S1: Within-host proviral half-life estimates from primary and sensitivity analyses**

| Participant | Analysis | Time to peak viral load (days) | Peak viral load (copies/ml) | Time to  setpoint viral load | Proviral DNA half-life (95% CI) in years |
| --- | --- | --- | --- | --- | --- |
| P1 | Primary^a^ | 31 | 53300 | 41 days | 0.41 (0 - 1.03) |
|  | Sensitivity 1^b^ | 23 | 12902 | 22 weeks | 0.41 (0 - 1.05) |
|  | Sensitivity 2^c^ | 14 | 1954 | 6 weeks | 0.41 (0 - 1.08) |
|  | Sensitivity 3^d^ | 30 | 71550 | 43 weeks | 0.33 (0 – 0.75) |
| P2 | Primary | 31 | 53300 | 41 days | 8.88 (0 – 77.36) |
|  | Sensitivity 1 | 23 | 12902 | 22 weeks | 34.6 (0 – 1115.49) |
|  | Sensitivity 2 | 14 | 1954 | 6 weeks | No decay |
|  | Sensitivity 3 | 30 | 71550 | 43 weeks | 2.3 (0 – 6.72) |
| P3 | Primary | 31 | 53300 | 41 days | 1.15 (0 – 2.86) |
|  | Sensitivity 1 | 23 | 12902 | 22 weeks | 1.4 (0 – 4.05) |
|  | Sensitivity 2 | 14 | 1954 | 6 weeks | 3.04 (0 – 18.08) |
|  | Sensitivity 3 | 30 | 71550 | 43 weeks | 0.9 (0 – 1.9) |
| P4 | Primary | 31 | 53300 | 41 days | 1.56 (0 – 8.66) |
|  | Sensitivity 1 | 23 | 12902 | 22 weeks | 1.56 (0 – 8.81) |
|  | Sensitivity 2 | 14 | 1954 | 6 weeks | 1.64 (0 – 9.85) |
|  | Sensitivity 3 | 30 | 71550 | 43 weeks | 1.4 (0 – 6.51) |

^a^ The primary analysis used peak viremia kinetics from a study of viremic controllers (reference 42 in manuscript )‬‬‬‬‬‬‬‬‬‬‬‬‬‬‬‬‬‬‬‬‬‬‬‬‬‬‬‬‬‬‬‬‬‬‬‬‬‬‬‬‬‬‬‬‬‬‬‬‬‬‬‬‬‬‬‬‬‬‬‬‬‬

^b, c, d^ The sensitivity analyses used median, minimum and maximum peak viremia kinetics from studies of elite controllers (references 43, 44 in manuscript)
