## Supplemental Table 2 for "HIV proviral burden, genetic diversity, and dynamics in viremic controllers who subsequently initiated suppressive antiretroviral therapy"

**Table S2**: Best-fit nucleotide substitution models

| Participant ID | Model ^a^ | LogL ^b^ | AIC ^c^ | w-AIC ^d^ |
| --- | --- | --- | --- | --- |
| P1 | GTR+F+I+G4 | -2056.7742 | 4459.5484 | 0.5537 |
|  | TVM+F+I+G4 | -2058.0462 | 4460.0924 | 0.4218 |
| P2 | K3Pu+F+I+G4 | -2779.8204 | 5951.6407 | 0.1185 |
|  | TPM2+F+I+G4 | -2779.8851 | 5951.7702 | 0.111 |
|  | TPM2u+F+I+G4 | -2779.8864 | 5951.7729 | 0.1109 |
|  | HKY+F+I+G4 | -2780.8967 | 5951.7933 | 0.1098 |
|  | TVM+F+I+G4 | -2778.0324 | 5952.0648 | 0.0958 |
|  | TIM+F+I+G4 | -2779.1122 | 5952.2244 | 0.0885 |
|  | TIM2+F+I+G4 | -2779.1412 | 5952.2825 | 0.0859 |
|  | TN+F+I+G4 | -2780.203 | 5952.406 | 0.0808 |
|  | GTR+F+I+G4 | -2777.3209 | 5952.6418 | 0.0718 |
| P3 | K3Pu+F+I+G4 | -1749.2209 | 3834.4418 | 0.3739 |
|  | TVM+F+I+G4 | -1747.8519 | 3835.7037 | 0.199 |
|  | TIM+F+I+G4 | -1749.1842 | 3836.3684 | 0.1427 |
|  | K3Pu+F+I | -1751.742 | 3837.484 | 0.0817 |
|  | GTR+F+I+G4 | -1747.851 | 3837.7019 | 0.0733 |
| P4 | TVM+F+I+G4 | -5522.3216 | 12200.6432 | 0.7346 |
|  | GTR+F+I+G4 | -5522.3398 | 12202.6795 | 0.2654 |

^a^ GTR = General Time Reversible model; TVM = Transversion model; K3Pu = Kimura 3-parameter model with unequal base frequencies; TPM2 = 3-parameter model with equal base frequencies; TPM2u = 3-parameter model with unequal base frequencies; HKY = Hasegawa, Kishino and Yano model; TIM = Transition model and unequal base frequencies; TIM2 = Transition model; TN = Tamura and Nei model. These models include empirical base frequencies (+F), invariable sites (+I), gamma distributed rate variation across sites with 4 rate categories (+G4).

^b^ LogL = Log likelihood

^c^ AIC = Akaike Information Criterion scores

^d^ w-AIC = Weighted AIC scores. Positive values denote nucleotide substitution models that were not significantly different from each other

^a^ GTR = General Time Reversible model; TVM = Transversion model; K3Pu = Kimura 3-parameter model with unequal base frequencies; TPM2 = 3-parameter model with equal base frequencies; TPM2u = 3-parameter model with unequal base frequencies; HKY = Hasegawa, Kishino and Yano model; TIM = Transition model and unequal base frequencies; TIM2 = Transition model; TN = Tamura and Nei model.

^b^ These models include empirical base frequencies (+F), invariable sites (+I), gamma distributed rate variation across sites with 4 rate categories (+G4).

^c^ LogL = Log likelihood

^d^ AIC = Akaike Information Criterion scores

^e^ w-AIC = Weighted AIC scores. Positive values denote nucleotide substitution models that were not significantly different from each other
